# Systematic dissection of epigenetic age acceleration in normal breast tissue reveals its link to estrogen signaling and cancer risk

**DOI:** 10.1101/2024.10.31.619735

**Authors:** Mary E. Sehl, Wenbin Guo, Collin Farrell, Natascia Marino, Jill E. Henry, Anna Maria Storniolo, Jeanette Papp, Jingyi Jessica Li, Steve Horvath, Matteo Pellegrini, Patricia A. Ganz

## Abstract

Breast aging encompasses intricate molecular and cellular changes that elevate cancer risk. Our study profiled DNA methylation and gene expression of 181 normal breast samples and systematically evaluated eight epigenetic clocks. We found that clocks trained using breast tissues demonstrate improved age prediction in normal breast tissue, and bias universally exists in epigenetic clocks, necessitating a proper definition of age acceleration. Cell composition analysis revealed significant age-related alterations and highlighted its distinct associations with age acceleration, including increased luminal epithelial and myoepithelial cells and reduced adipocytes and immune cells, connecting age acceleration to carcinogenesis from a cell compositional perspective. Additionally, CpG sites associated with age acceleration were enriched for estrogen receptor binding sites, providing a mechanistic link between estrogen exposure, accelerated aging, and cancer. These findings highlight the importance of cellular heterogeneity in epigenetic age estimates and the potential of age acceleration to guide for risk stratification and prevention strategies.

## Introduction

Aging is an intricate biological process accompanied by progressive molecular, cellular, and tissue-level changes, leading to functional decline and increased disease susceptibility^1^. In breast tissue, aging induces a cascade of molecular alterations such as DNA damage accumulation^2^, telomere shortening^3^ and epigenetic modifications^4^. At the cellular level, these changes manifest as senescence^5^, altered cell composition and disrupted tissue architecture^6^. Specifically, breast aging is associated with the accumulation of dysfunctional luminal epithelial cells^7^, a reduction in hormone-sensitive cells^8^, and an increase in immune cell infiltration along with a decline in adaptive immune cells^9^. These alternations contribute to genome instability and a pro-inflammatory environment with compromised immune surveillance, heightening the risk of malignant transformation and breast cancer development. Understanding these aging-related changes is essential for developing strategies to alleviate the adverse effects of aging on breast health and prevent breast cancer.

The concept of the epigenetic clock has emerged as a powerful tool for studying aging and elucidating its molecular mechanisms. Various epigenetic clocks have been developed over the years^10–16^, each leveraging DNA methylation patterns at specific genomic loci to accurately measure biological age. Previous research has shown significant associations between epigenetic age and a wide range of age-related diseases, including breast cancer^17^. Beyond reflecting biological age, deviations in epigenetic age from a normal trajectory—known as age acceleration or deceleration— have garnered considerable attention due to their profound implications for diverse health outcomes^18^. In the breast aging field, pioneering work has established the link between accelerated epigenetic aging with lifetime estrogen exposure^19^ and increased breast cancer risk^20^. Despite these advancements, a comprehensive understanding of breast age acceleration, particularly its mechanistic link with estrogen exposure and cancer risk from molecular and cellular perspectives, remains elusive.

A critical gap in current research is the lack of consideration for tissue heterogeneity. Breast tissue consists of diverse cell types, each potentially aging at different rates and in distinct ways, while most existing epigenetic clocks are built with bulk DNA methylation mainly using blood samples^21^, failing to account for the complexity and variability of breast tissue. This raises questions about the accuracy of these clocks in predicting breast biological age and poses additional challenges on understanding breast-specific aging processes. Additionally, most epigenetic aging studies overlook the complex cell compositional changes with age^21^, focusing instead on bulk tissue analysis, which can obscure insights into the dynamics of specific cell types, their distinct relationships with epigenetic age and age acceleration, and their implications for cancer risks. Detailed investigations and characterizations of breast age acceleration from cellular, epigenetic, and transcriptomic perspectives are needed to elucidate breast aging processes, providing insights into potential intervention targets to promote breast health.

To address these gaps, we profiled and analyzed the DNA methylation and gene expression of 181 normal breast samples aged 19 to 90 (Figure 1A, 1B). By deconvolving the cell type abundance of breast tissue, we comprehensively studied the epigenetic aging at both the molecular and cellular levels. First, we systematically evaluated the predictive accuracy of eight epigenetic clocks in normal breast tissue, including two pan-tissue, two breast-specific, two second-generation, and two first-generation clocks. Our findings show that pan-tissue and breast-specific clocks accurately predicted age, while blood-based clocks underperformed due to tissue heterogeneity. We also revealed systematic biases in epigenetic clocks for age prediction and advocate for a proper definition of age acceleration to avoid confounding by chronological age. Cell type abundance analysis indicated significant age-related changes in breast tissue composition, including increased adipocytes and vascular endothelial cells, and decreased luminal epithelial and basal myoepithelial cells. Notably, epigenetic age acceleration was associated with distinct cellular changes, suggesting a higher risk of breast carcinogenesis from a cellular population perspective. We identified CpG sites associated with age acceleration, enriched for estrogen receptor binding sites (ESR1), linking estrogen exposure to accelerated breast aging and increased cancer risk from the molecular level. Transcriptome analysis revealed differentially expressed genes associated with age acceleration for each clock, while the overlap among different clocks is minimal and the pathway enrichments reflect unique biological signals captured by different clocks. Overall, our findings emphasize the need for considering cellular and tissue heterogeneity in epigenetic aging studies, providing valuable insights into the molecular mechanisms linking estrogen exposure, epigenetic aging, and breast cancer risk. Future research should focus on refining epigenetic clocks and exploring their clinical applications in breast aging and cancer risk assessment.

**Figure 1.**
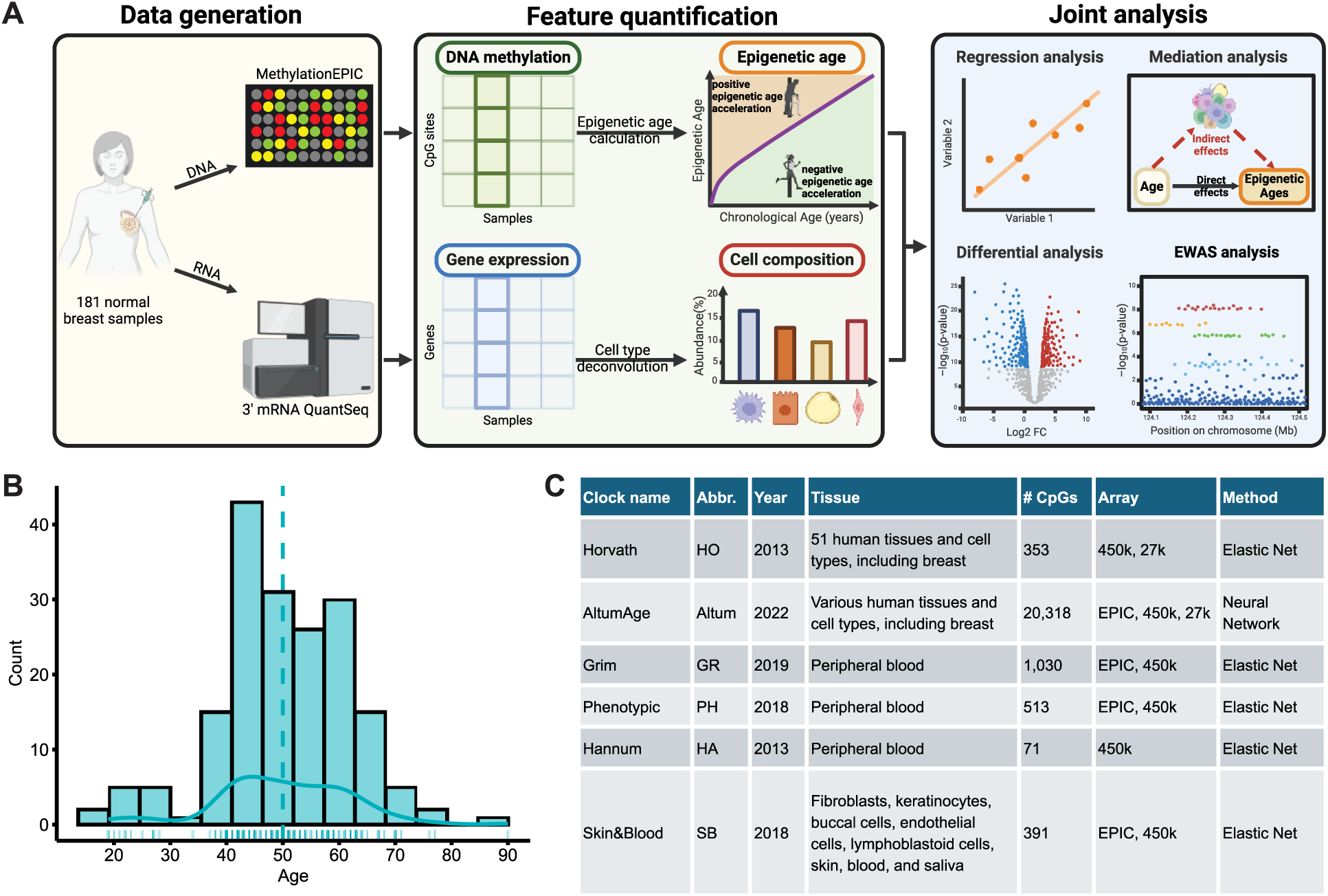
Study design and overview. **(A)** 181 normal breast tissue samples from Komen Tissue Bank (KTB) with paired DNA methylation and gene expression data were analyzed in this study. DNA methylation data were used to calculate the epigenetic age for different clocks. Gene expression data were used to estimate cell type abundances. The DNA methylation, gene expression, epigenetic age, and cell composition were then jointly analyzed to comprehensively characterize the epigenetic aging and cell compositional landscape of normal breast tissue during aging. **(B)** Histogram and density plot showing chronological age distribution of the 181 study samples. The vertical dashed line shows the median age (50 years old). The solid line represents the distribution density. **(C)** Table summarizing the features of six popular clocks including two pan-tissue clocks: Horvath clock and AltumAge, two second-generation clocks: GrimAge and Phenotypic Age, two first-generation clocks: Hannum clock and Skin&Blood clock. Besides the six published clocks, we also trained two breast-specific clocks in KTB samples using Elastic Net algorithms and Epigenetic Pacemaker and included them in following evaluations.

## Results

### Age prediction accuracy of epigenetic clocks in normal breast tissue

Evaluating the age prediction accuracy in healthy breast tissue using eight epigenetic clocks (Figure 1C, Methods), we observed strong correlations between epigenetic age and chronological age across most clocks, except for Phenotypic Age (Figure 2, Table 2). Significant correlations among different epigenetic age estimates were also noted (Supplementary Figure 3A). Notably, the pan-tissue and breast-specific clocks demonstrated high accuracy in predicting age in breast tissue, with the ElasticNet clock trained on Komen Tissue Bank (KTB) breast tissue showing the highest accuracy (Pearson Correlation Coefficient (Corr.) = 0.94, mean absolute error (MAE) = 3.33 years). Other clocks such as HO (Corr. = 0.90, MAE = 7.92 years), Altum (Corr. = 0.88, MAE = 5.24 years), and EPM (Corr. = 0.78, MAE = 6.56 years) also performed well.

**Figure 2.**
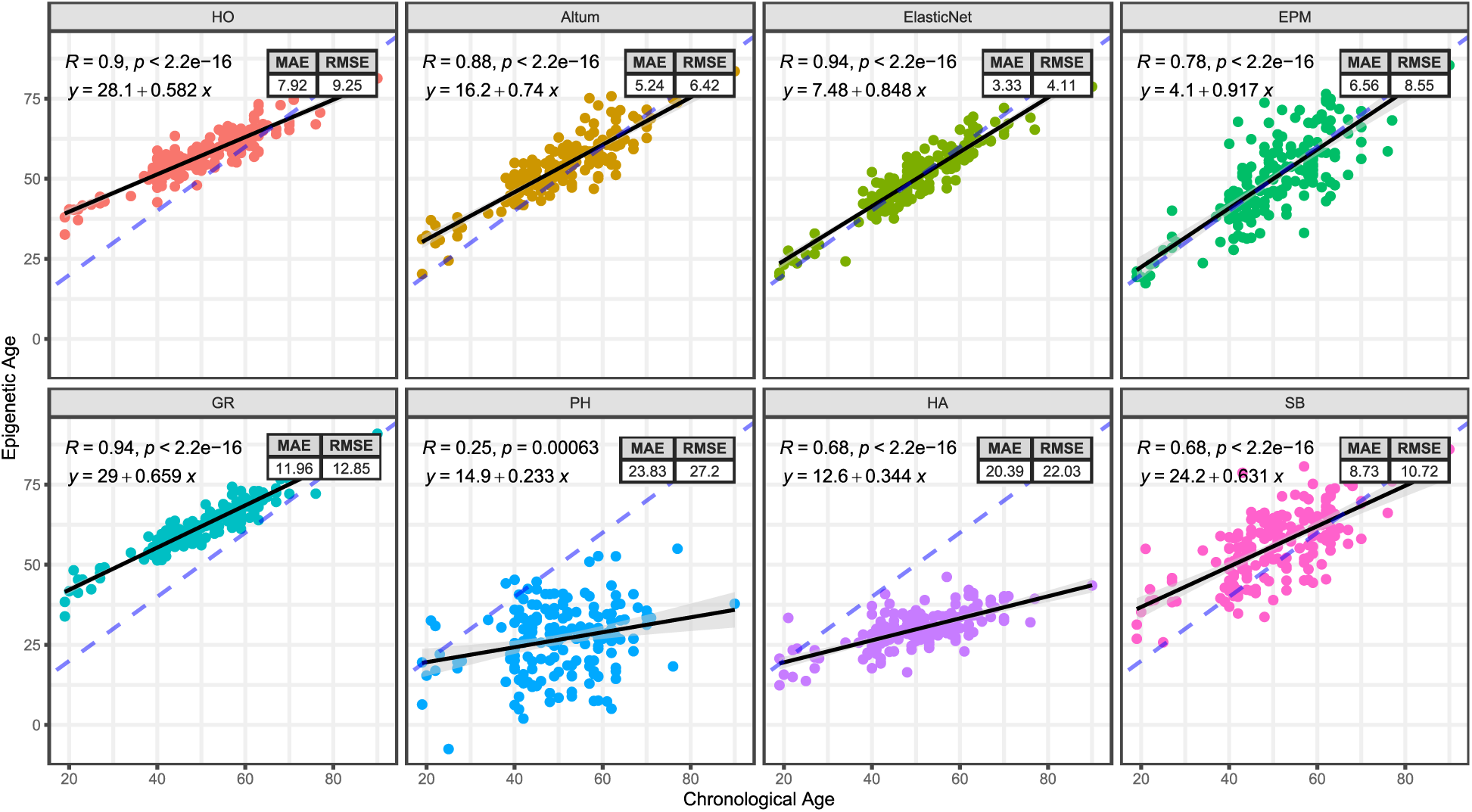
Age prediction accuracy across eight epigenetic clocks. Top panel shows the epigenetic clocks for pan-tissue (HO: Horvath clock, Altum: AltumAge) and breast-specific clocks (ElasticNet: clock trained with Elastic Net algorithm, EPM: clock trained with Epigenetic Pacemaker). The bottom panel shows blood-based epigenetic clocks including second-generation clocks (GR: GrimAge, PH: Phenotypic Age) and first-generation clocks (HA: Hannum clock, SB: Skin&Blood clock). The dashed blue line shows *y* = *x* diagonal line, black line represents the regression line between epigenetic age and chronological age for each clock, with shadowed region indicating 95% Confidence Interval for regression. Pearson correlation coefficient statistics, regression line equation, and prediction error (MAE: mean square error, RMSE: root mean square error) are annotated to the top of each panel on the plot.

Conversely, blood-based (first- and second-generation) clocks that did not include breast tissue in their original training set showed significantly lower correlations or higher MAEs, indicating reduced predictive accuracy. An exception was GrimAge, which includes chronological age as a predictor and thus showed a high correlation of Corr.= 0.94, similar to the best-performing breast-specific clocks, but with a much higher MAE (11.96 years). These findings suggest that blood-based clocks tend to underperform for predicting age in breast tissue, likely due to tissue heterogeneity. This underscores the importance of using tissue-specific or pan-tissue clocks to minimize cross-tissue biases and improve age prediction accuracy.

Our analysis also revealed that the regression line between predicted (epigenetic) age and chronological age consistently exhibited a positive intercept and a slope less than 1, regardless of the clock used. The large positive intercepts of first- and second-generation clocks, indicating predicted age in breast tissue at chronological age zero, might suggest that breast tissue appears “older” compared to other tissues such as blood^22^. However, the slope being less than 1 raises concerns about underestimating age in older subjects, highlighting a systematic bias in the epigenetic clock models. This issue will be further discussed in the following section.

### Systematic biases in epigenetic clocks and justification of age acceleration

To elucidate the observed systematic biases in age estimation stemming from model bias, we simulated a scenario where DNA methylation can fully explain the variance in chronological age. Using penalized regression techniques (LASSO, Ridge, and Elastic Net regression) to construct the clocks, we consistently found that the regression slope of predicted age on chronological age was below 1 (Supplementary Figure 1A). This occurs because penalized regression methods, used to manage the high-dimensional challenge where features (CpG sites) outnumber samples, shrink regression coefficients towards zero. As a result, predicted age (epigenetic age) rarely matches chronological age exactly and tends to have reduced variance compared to chronological age. Consequently, the prediction error (predicted age - chronological age) correlates with chronological age (Supplementary Figure 1C).

These biases complicate the definition of age acceleration. Throughout the literature, we recognized various studies have used different definitions, with no consensus reached. Some studies^10,16,21,23–25^ define the age acceleration as the difference between epigenetic age and chronological age (difference definition), while others^13,14,26–29^ define it as the deviation of epigenetic age from its expected value given chronological age (residual definition). A slope smaller than 1 indicates that age difference is inversely associated with chronological age (Supplementary Figure 1C, 2C), leading to spurious associations between health outcomes and age differences due to confounding by chronological age. In contrast, the residual definition represents the part of epigenetic age unexplained by chronological age and is orthogonal to chronological age (Supplementary Figure 1B, 2B), eliminating age confounding concerns. In this study, we advocate against the use of age difference definition since it can easily lead to false discoveries when not calibrated with chronological age; And we defined the age acceleration as the regression residual between epigenetic age and chronological age.

### Age and other demographic related changes in breast cell composition

We next investigated how cell type abundance changes with advancing age in breast tissue. Using CIBERSORTx^30^ with GTEx normal breast snRNA-seq data^31^ as a reference, we deconvolved the cell type abundance from bulk gene expression data for each sample and observed significant age-related changes in breast tissue composition (Figure 3A, 3B). Specifically, advancing chronological age is associated with a notable increase in the proportion of adipocytes (p<4.12e-8, adjusted p<1.32e-06) and vascular endothelial cells (p<1.02e-4, adjusted p<1.21e-03), and a significant decrease in proportions of luminal epithelial cells (p<9.1e-11, adjusted p<5.83e-09) and basal myoepithelial cells (p<1.13e-4, adjusted p<1.21e-03). Additionally, the proportion of immune dendritic cells/macrophages significantly increased (p<1.93e-6, adjusted p<4.11e-05), indicating an elevated inflammatory landscape with aging.

**Figure 3.**
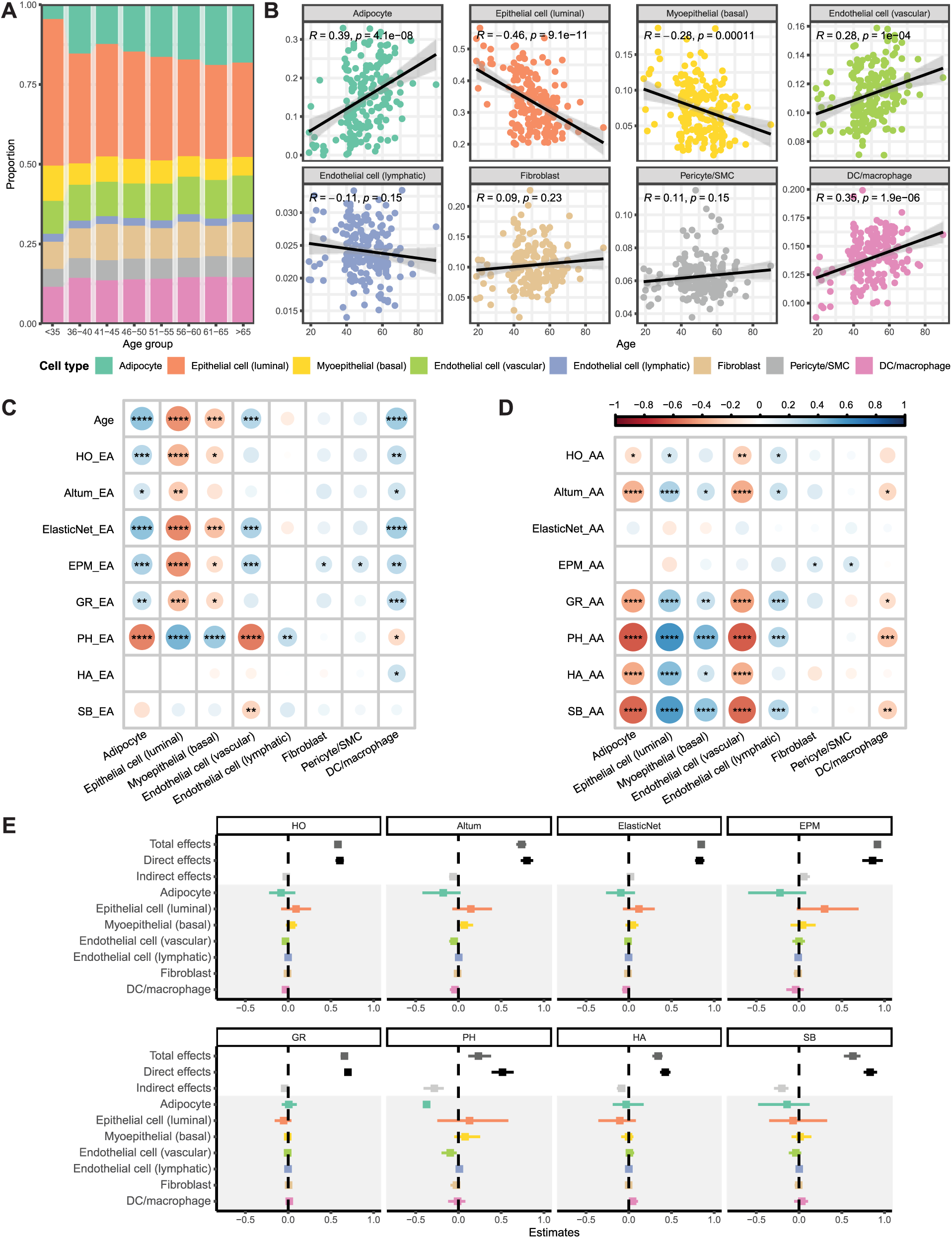
Cell composition’s correlation with epigenetic age and age acceleration. **(A**-**B**) Cell composition’s correlation with advancing chronological age**. (A**) Bar plot showing the average cell proportion dynamics across age groups. (**B**) Scatterplot showing the trend and strength of proportion change with chronological age for each cell type. Black line indicates the regression line between proportion and chronological age, with shadow region indicating 95% regression Confidence Intervals. The Pearson Correlation Coefficients and significance level are annotated to the top of each panel on the plot. **(C**-**D**) Heatmap showing the correlation between cell composition and epigenetic age (**C**) and age acceleration (**D**). Chronological age was added to the top row as a reference. Circle size and color scale representing the strength of correlation with blue color shows positive correlation and red color indicates negative correlation. Asterisk indicates significance level after p-value adjustment. *: adjusted p-value < 0.05; **: adjusted p-value < 0.01; ***: adjusted p-value < 0.001; ****: adjusted p-value < 0.0001. **(E**) Forest plot displaying the cell composition’s mediation effects on the relationship from chronological age to epigenetic age for each clock. The total effects, direct effects and indirect effects are annotated on the top. Grey shadowed region shows the individual contribution of each cell type to epigenetic age. The dot represents the effects from mediation analysis, with the width of the line representing the 95% Confidence Interval.

We also examined the associations between cell type abundance and various demographic and reproductive variables (Supplementary Data 3). Hispanic ethnicity was associated with a lower proportion of fibroblasts (p<0.017). Higher body mass index (BMI) correlated with increasing proportions of immune dendritic cells/macrophages (p<0.014) and decreased luminal epithelial cells (p<0.033). Tobacco smoking history was linked to a reduced proportion of pericytes/smooth muscle cells (p<0.034), though these associations did not remain significant after p-value adjustment. Interestingly, adipocyte proportion did not correlate with BMI (Supplementary Figure 6), suggesting that an individual BMI is primarily related to overall adiposity rather than the proportion of adipocytes in breast tissue. There were no significant associations between cell proportions and age at menarche, parity, or history of breastfeeding.

### Changes in breast cell composition with breast epigenetic age

Given that current epigenetic clocks primarily focus on bulk tissues with surrogate measures of DNA methylation, it is intuitive that changes in cell composition could influence epigenetic age. We investigated how cell type proportions correlate with epigenetic age for each clock, using chronological age as a reference (Figure 3C). Our analysis revealed significant associations between epigenetic age estimates and at least one cell type proportion for almost all clocks. The correlation patterns generally mirrored those observed with chronological age, except for the Phenotypic Age, Hannum clock, and Skin&Blood clock, which did not predict age accurately in breast tissue.

Specifically, adipocyte proportions displayed negative correlations with epigenetic ages across multiple clocks. Vascular endothelial cells consistently showed positive correlations with epigenetic age, indicating vascular remodeling. Luminal epithelial and basal myoepithelial cells exhibited strong negative correlations, reflecting a decline in breast density with epigenetic aging. The positive correlation between immune dendritic cells/macrophages and epigenetic age highlights an increased inflammatory environment. These findings underscore the complexity of epigenetic aging, validating the biological relevance of epigenetic clocks in capturing aging-related cellular changes, and emphasize the importance of considering these changes when interpreting epigenetic age estimates.

Beyond the major cell types, we also examined the correlation between epigenetic age and immune cell enrichment scores calculated by immuneCellAI^32^ using gene expression data. We found that macrophage scores positively correlated with both chronological and epigenetic age, while gamma-delta T (γδT) cell scores displayed a negative correlation (Supplementary Figure 5A), both well aligning with previous observations^9,33^. The increased macrophage enrichment scores highlight an elevated inflammatory status, while decreased γδT cell enrichment scores potentially reflect a decline in maintenance of tissue homeostasis^33^. Together, these findings indicate altered immune surveillance dynamics during aging in normal breast microenvironment.

### Changes in breast cell composition with breast age acceleration

We next investigated whether epigenetic age accelerations are associated with cell compositional changes in breast tissue. Figure 3D illustrates the correlations between epigenetic age acceleration and cell abundance for each clock measure and cell type. We first noted the correlation variability among different clocks: Breast-specific clocks, optimized for predicting chronological age, showed the least correlation with cell abundance. In contrast, other clocks demonstrated significant associations with cell compositional changes. Specifically, blood-based clocks (both first- and second-generation) showed significant correlations with immune cell enrichment scores after p-value adjustment (Supplementary Figure 5B). Notably, the pan-tissue clock AltumAge exhibited greater sensitivity to cellular changes compared to the Horvath clock, likely due to it used higher number (70 times more) of CpG sites for age prediction.

At the cell-type level, interestingly, we observed that adipocyte and vascular endothelial cell proportions negatively correlated with epigenetic age acceleration across pan-tissue, first-, and second-generation clocks (p<0.01, adjusted p-value< 0.05). This suggests a decrease in these cells with advancing epigenetic age, contrasting with their positive correlation with chronological age. Conversely, Luminal epithelial cells, basal myoepithelial cells lymphatic endothelial cells showed significant positive correlations with age acceleration across clocks. Additionally, the proportion of immune dendritic cells/macrophages was significantly lower in accelerated breast aging tissue, reflecting compromised immune surveillance. These findings underscore the distinct characteristics of accelerated breast aging compared to normal aging. The increase in luminal epithelial and basal myoepithelial cells, along with the reduction in immune cells in accelerated aging breast, suggests a potential higher risk of breast carcinogenesis from a cell compositional perspective.

To further explore whether cell composition mediates the increase in epigenetic age during aging process, we conducted a mediation analysis. Consistent with the correlation analysis findings, we found significant mediation effects for blood-based clocks (first- and second-generation), with GrimAge being an exception due to its inclusion of chronological age in the predictors. Our analysis also revealed that adipocytes negatively contribute to epigenetic age, while luminal epithelial and basal myoepithelial cells positively contribute, although the strength of these mediating effects were marginally significant.

### Association between breast age acceleration and breast cancer risk measurement

We further examined the association between breast epigenetic age acceleration and breast cancer risk using the Breast Cancer Risk Assessment Tool (Gail Model) and the Tyrer-Cuzick Risk Calculator (IBISv8, for 10-year and lifetime breast cancer risk). Supplementary Figure 7A presents the correlations between epigenetic age acceleration and breast cancer risk estimates for each risk score and clock. Among them, only Horvath clock demonstrated a marginally significant association (corr.=0.14, p=0.054) and lifetime Tyrer-Cuzick score (corr.=0.14, p=0.068). Interestingly, after adjusting for the cell proportions, we observed an increase in correlation between Horvath age acceleration and 10-year Tyrer-Cuzick score (corr.=0.17, p=0.022) and lifetime Tyrer-Cuzick score (corr.=0.17, p=0.022) (Figure 4A), while the correlation for other clocks remains insignificant (Supplementary Figure 7B).

**Figure 4.**
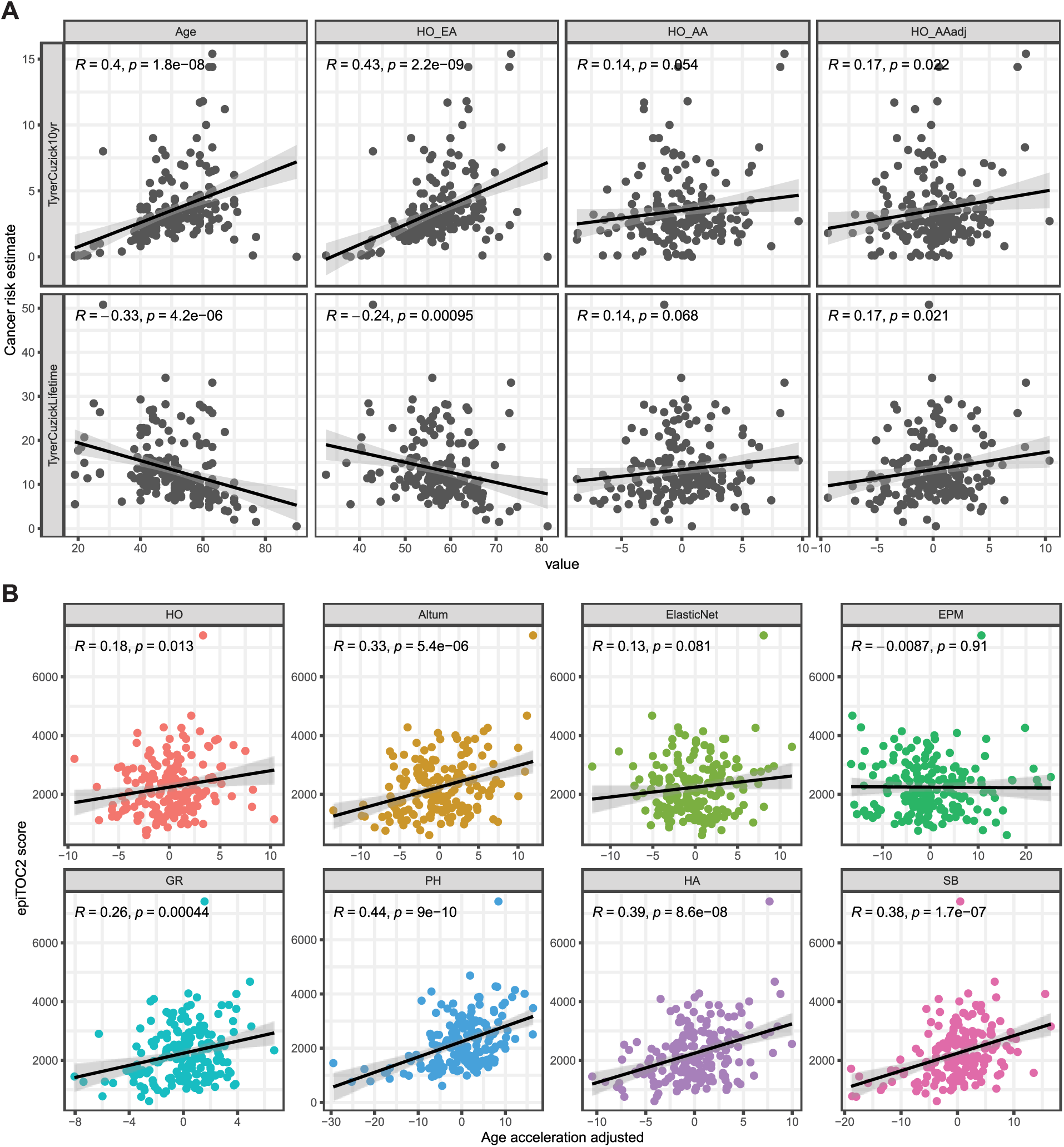
Age acceleration’s association with cancer risk estimates. (**A**) Scatter plot showing the association between Tyrer-Cuzick score and chronological age, Horvath clock’s epigenetic age, age acceleration, and age acceleration adjusted by cell proportions (top panel: 10-year risk, bottom panel: lifetime risk). (**B**) Scatter plot showing the association between DNAm based cancer risk (epiTOC2 score) and age acceleration adjusted by cell proportions for each clock. Black line represents the regression line with shadowed region indicating 95% Confidence Intervals. Pearson Correlation Coefficients and significance level are annotated to the top of each panel on the plot.

Beyond risk estimations derived from demographic and reproductive history, we also assessed DNA methylation-based risk using epiTOC2^34^, a breast cancer risk calculator which quantifies the total number of stem cell divisions. All but breast-specific clocks displayed a significant positive association between epiTOC2 score and age acceleration adjusted by cell proportions. These results showed that the link between age acceleration and cancer risk is not purely due to the cell compositional changes, although different clocks showed varying association strengths with cancer risk scores.

### Identification of CpG sites associated with breast age acceleration and its link to estrogen receptor

To characterize the CpG sites associated with age acceleration after adjusting for cell proportions, we performed an epigenome-wide association study (EWAS). Supplementary Figure 8 presents the QQ and Manhattan plot of the EWAS analysis results for each clock. CpG sites with q-value smaller than 0.05 are defined as significant and summarized in Supplementary Data 5 for each clock, along with their nearby genes annotated. Notably, blood-based clocks identified more significant CpG sites associated with age acceleration compared to both pan-tissue clocks and breast-specific clocks.

To better understand the biological mechanisms underlying these CpG sites, we performed genomic region set enrichment analysis on the significant CpG sites, using the total CpG sites as the background. The top 10 most significantly enriched Transcription Factor Binding Sites (TFBS) of each clock were collected and visualized in Figure 5. Our analysis revealed that estrogen receptor 1 (ESR1) binding sites were among the top 10 enriched TFBS for most clocks examined, providing a potential link between age acceleration and estrogen exposure. Although the association between age acceleration and estrogen exposure has been reported in our previous study^19^, this analysis provides the direct molecular evidence supporting the association for the first time.

**Figure 5.**
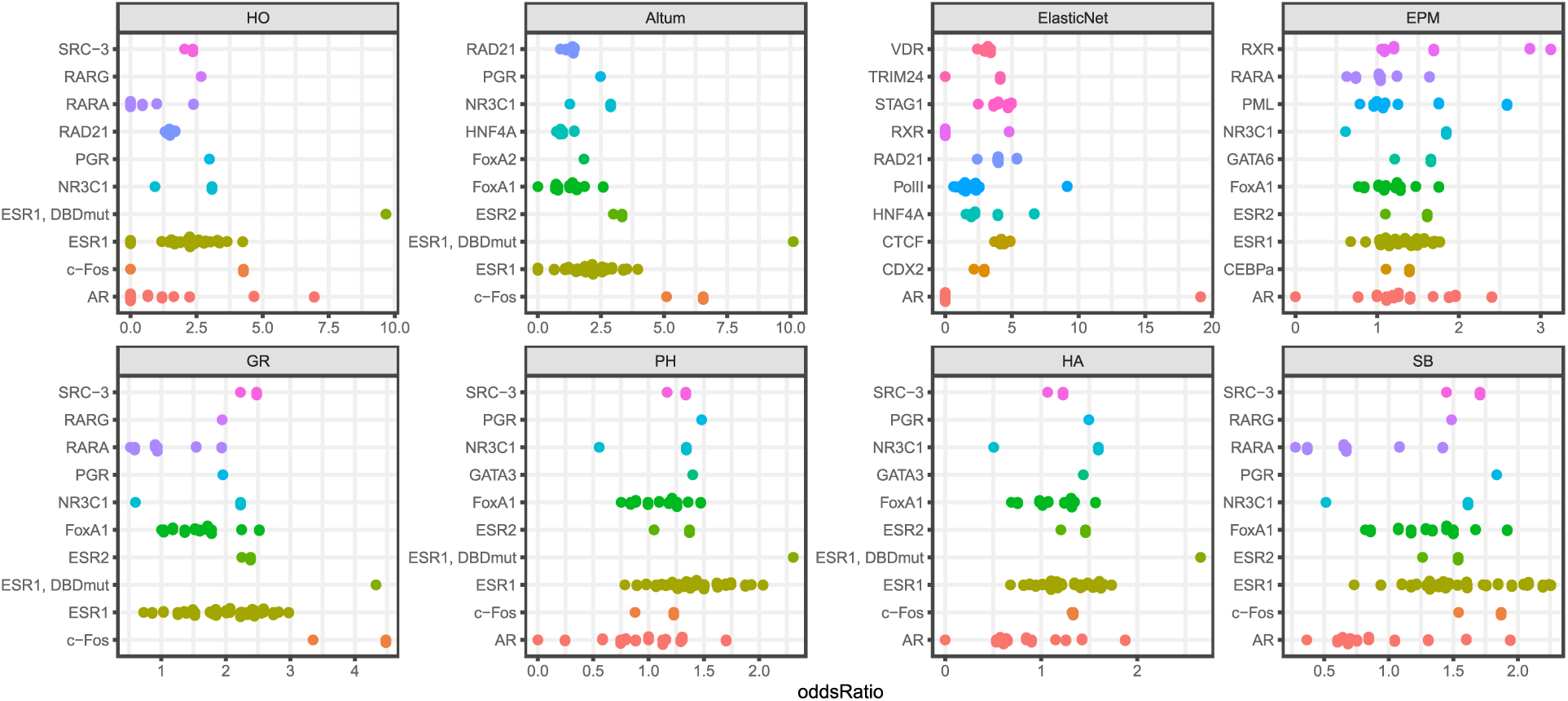
Age-acceleration-related CpG sites enriching for estrogen receptor binding sites. EWAS analysis was performed to identify CpG sites associated with age acceleration after adjusting cell proportions for each clock, followed by genomic region enrichment analysis. The top 10 most significantly enriched Transcription Factor Binding Sites (TFBS) for each clock were visualized, with dots indicating the enrichment odds ratio of genomic regions for a particular TFBS. Estrogen receptor 1 (ESR1) binding sites were found among the top 10 most significantly enriched TFBS for most clocks examined.

Estrogen is known to stimulate the division and proliferation of breast tissue, leading to increased cellular turnover and DNA double-strand breaks^35^. This heightened cellular activity, coupled with the accumulation of DNA damage, promotes genomic instability—a hallmark of cancer development. Recent research also found that DNA double-strand breaks erode the epigenetic landscape, contributing to mammalian aging^36^. Thus, the enrichment of ESR1 binding sites for age acceleration-associated CpG sites provides a plausible mechanism linking age acceleration and increased cancer risk: estrogen exposure accelerates aging through DNA double-strand breaks. These findings suggest that estrogen exposure may have a significant role in the epigenetic changes observed in accelerated breast aging, potentially heightening breast cancer risk.

### Transcriptomic alternations associated with breast age acceleration

To identify genes whose expression is associated with age acceleration, we conducted a differential gene expression (DE) analysis using a regression framework, testing the associations between each gene’s expression and age acceleration, with chronological age included as a covariate. This analysis was conducted for each epigenetic clock, both before and after adjusting for cell proportions. We found that while thousands of genes were identified as DE genes before accounting for cell compositional changes, this number dramatically decreased after adjusting for cell proportions (Supplementary Figure 9A), indicating that many observed differences in gene expression were primarily due to cell population dynamics. Focusing on the DE genes after adjusting cell proportions, we noticed there was minimal overlap among different clocks (Figure 6A), suggesting that each clock captures distinct biological signals.

**Figure 6.**
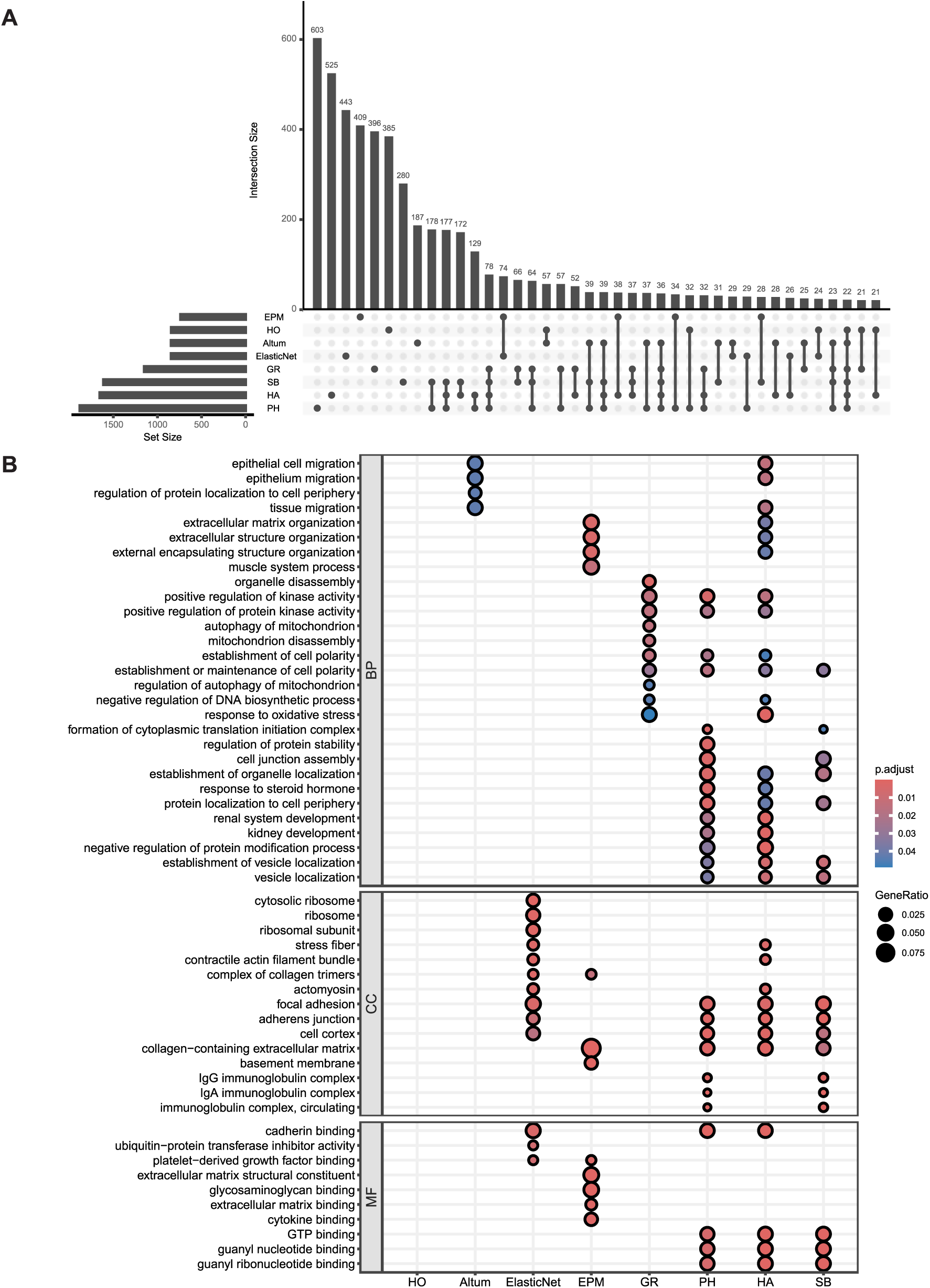
Transcriptome analysis characterizing genes and pathways associated with age acceleration. **(A)** Differential expression (DE) analysis was performed to identify genes significantly associated with age acceleration after adjusting cell proportions. Bar plot shows the set size of DE genes of each clock and their intersections. Little overlap was found on the DE genes among clocks. **(B)** Gene ontology (GO) enrichment analysis was performed on the DE genes of each clock to characterize the enriched pathways for each clock. Top 10 most significantly enriched pathways were shown for each clock and arranged into three categories (BP: Biological Pathways, CC: Cellular Component; MF: Molecular Function). Dot size represents the gene ratio of a specific pathway overlapping with DE genes and color scale represent the adjusted significance level of enrichment.

We also performed Gene Ontology (GO) enrichment analysis with the DE genes and fount the enrichment results revealed distinct patterns in biological pathways for each epigenetic clock (Figure 6B). Specifically, there were no enriched biological processes for the Horvath and ElasticNet clocks. While the Altum clock showed enrichment for processes related to epithelial migration, and the EPM clock highlighted pathways involved in extracellular matrix organization. Blood-based clocks demonstrated greater overlap in enriched biological pathways, emphasizing processes such as kinas activity regulation, steroid hormone signaling and cell polarity. Furthermore, Gene Set Enrichment Analysis (GSEA) based on the DE analysis shows a similar pattern where different clocks enrich for diverse pathways, with the blood-based clocks to have bigger overlap, featuring immune and metabolic processes. (Supplementary Figure 9B).

Collectively, these findings in the transcriptome analysis underscore the critical importance of accounting for cell composition in epigenetic studies^37^. The pathway enrichment analysis also highlight that different epigenetic clocks capture unique and complementary aspects of biological aging in breast tissue, providing a nuanced understanding of the underlying molecular mechanisms. This insight highlights the complexity of breast aging and interplay between epigenetic dynamics and transcriptomic variations in breast aging processes, and further investigation are needed to advance the precision and applicability of epigenetic clocks in breast aging research and clinical interventions.

## Discussion

In this study, we systematically evaluated the predictive accuracy of eight epigenetic clocks in normal breast tissue, the associations between epigenetic age estimates and cell composition, and their potential implications for breast cancer risk. Our findings demonstrate that pan-tissue and breast-specific epigenetic clocks exhibit superior accuracy in age prediction compared to clocks developed for other tissues. This discrepancy underscores the limitations of cross-tissue applications of epigenetic clocks and emphasizes the importance of considering tissue specificity in epigenetic age estimation. Our analysis also revealed systematic biases in age prediction, with regression lines consistently showing positive intercepts and slopes less than one. Simulations indicated that these biases arise from the shrinkage of regression coefficients in penalized regression, which is common in high-dimensional data settings (e.g., epigenetic clock). While previous studies have reported the underestimate bias in the old subjects of Horvath clock^38^, we are, to our best knowledge, the first to illustrate that the bias is a universal problem for all epigenetic clocks examined, and that it is rooted in the statistical models used to construct the clocks. These biases complicate the definition of age acceleration. To address this, we advocate for the residual definition of age acceleration, which eliminates confounding by chronological age and provides a more robust measure for studying age-acceleration-related health outcomes.

Using cell type abundance estimated from transcriptomic data, we confirmed several cell compositional changes associated with advancing chronologic age described in previous studies, including increased proportions of adipocytes and vascular endothelial cells^39^, and decreases in luminal epithelial and basal myoepithelial cells^40^. We also found that immune dendritic cells/macrophages increased with advancing chronological age in healthy breast tissue. These findings indicate an age-associated remodeling of breast tissue, characterized by a decline of breast density and an elevated inflammatory landscape. The observed correlations between cell composition and epigenetic age mirrored those with chronological age, validating the biological relevance of epigenetic clocks in capturing aging-related cellular alterations. However, the variability of correlations also highlights the influence of cell composition on epigenetic/biological age estimates, emphasizing the necessity of considering cell composition heterogeneity when interpreting epigenetic aging data.

While previous studies have examined cell compositional changes in breast tissue with chronologic age, this is the first study to characterize the association between cell composition and epigenetic age accelerations in normal breast tissue. Interestingly, we found that epigenetic age acceleration of multiple clocks is associated with distinct changes of cell composition, with a significant increase in both luminal epithelial and basal myoepithelial cell proportions, and a decrease in adipocytes, vascular endothelial cells and immune dendritic cell/macrophages. The rise in luminal and basal myoepithelial cells may heighten breast cancer risk, as these cells are progenitors of common breast cancer subtypes^41^. Reduced adipocytes and vascular endothelial cells can disrupt hormonal balance^42^ and angiogenesis^43^ respectively, potentially affecting tissue homeostasis and contributing to an environment that favors tumor development. Lastly, the decrease in immune dendritic cells/macrophages suggests a compromised immune surveillance^9^, allowing abnormal cells to proliferate unchecked. These changes jointly highlight the unique cellular dynamics of accelerated breast aging and implies a higher potential for breast carcinogenesis from a cellular population perspective.

Our study also explored the associations between breast epigenetic age acceleration and breast cancer risk. We assessed the association between epigenetic age acceleration and Gail or Tyrer-Cuzick scores and did not find significant association. Interestingly, age acceleration in the Horvath clock has a significant association with Tyrer-Cuzick score after adjusting for cell proportions, suggesting that cellular heterogeneity plays a role in modulating cancer risk. The positive correlation between age acceleration and total number of stem cell divisions, as measured by epiTOC2, supports the link between accelerated epigenetic aging and increased cancer risk. These findings highlight the potential utility of epigenetic clocks in assessing breast cancer risk and underscore the importance of considering cell composition in these cancer risk analyses.

Epigenome-wide association studies (EWAS) have identified a handful of CpG sites associated with age acceleration after adjusting for cell proportions, with significant enrichment for estrogen receptor 1 (ESR1) binding sites across multiple epigenetic clocks. Estrogen exposure has been found to promote DNA double-strand breaks^44^, which contribute to mammalian aging^36^, suggesting a mechanistic link between estrogen exposure and accelerated epigenetic aging. Estrogen-driven cellular proliferation and DNA damage induce genomic instability, a hallmark of both aging and cancer development^45,46^, linking the age acceleration with cancer risk. Although previous studies have reported age acceleration is potentially associated with estrogen exposure^19^ and breast cancer risk^47^, our findings provide the direct molecular evidence that estrogen exposure accelerates breast tissue aging, potentially heightening the risk of breast cancer. Complementing these epigenetic insights, differential gene expression analysis has revealed significant transcriptomic alterations linked to age acceleration, many of which are driven by underlying shifts in cell composition. Gene ontology and gene set enrichment analyses have highlighted distinct biological processes associated with different epigenetic clocks, reflecting their unique biological signals. These insights emphasize the complexity of breast aging, driven by the interplay between estrogen exposure, epigenetic dynamics, and transcriptomic variations during aging process. Understanding these interactions is crucial for refining epigenetic clocks and enhancing their applicability in clinical settings to assess aging and cancer risk.

There are some limitations of our work. Firstly, the cross-sectional nature of the data, similar to most epigenetic aging studies, makes it difficult to disentangle subject-specific effects and aging effects on the epigenetic landscape. We believe longitudinal data benefit us and profile the aging rates and trajectory more accurately. Secondly, the relatively short duration after sample collection did not allow us to assess breast cancer incidence among the study population; we relied on breast cancer risk scores like the Gail or Tyrer-Cuzick scores, which may not accurately reflect the future likelihood of developing cancer^48^. This reliance limits our ability to access the relationship between age acceleration and the true potential of developing breast cancer. Additionally, our analysis was based on bulk DNA methylation and gene expression data, which lacks single-cell level resolution. While we recovered some cell type-level dynamics via deconvolution, it can be further improved through the lens of single-cell techniques. We envision that, single-cell DNA methylation, coupled with single-cell transcriptomic profiling technologies, would provide a more detailed understanding of how epigenomic changes interplay with transcriptomics during the normal aging process. Future research should focus on longitudinal studies and single cell analyses to refine these clocks and explore their clinical applications in breast aging and cancer risk assessment.

In summary, our study underscores the critical need for tissue-specific epigenetic clocks to accurately predict age and assess age acceleration. The observed biases in age prediction models highlight the importance of using appropriate definitions for age acceleration. Age-related changes in breast cell composition and their impact on epigenetic age emphasize the need to consider cellular heterogeneity in aging studies and age acceleration shows distinct association with cell composition, suggesting a its link with cancer risk from the cell compositional perspective. Our findings also provide valuable insights into the molecular mechanisms linking estrogen exposure, epigenetic aging, and breast cancer risk, paving the way for future research and clinical interventions in breast aging. Future studies should focus on refining epigenetic clocks for broader applicability and investigating their potential for early detection and prevention of age-related diseases, including breast cancer.

## Methods

### Study samples and specimens

We utilized breast tissue specimens from the Susan G. Komen Tissue Bank (KTB) at the Indiana University Simon Comprehensive Cancer Center, a unique repository of samples from healthy female donors. Each tissue sample is well annotated with the donor’s race and ethnicity, height, weight, family history, reproductive history, and medication use. All participants in the study have provided informed consent and the study received approval from the UCLA Institutional Review Board. Data for this study were collected as part of a cross-sectional study designed to investigate the associations between breast epigenetic age and hormonal factors. We initially recruited 200 female participants, categorized into four groups: (i) premenopausal and nulliparous, (ii) premenopausal and with at least 1live birth, (iii) postmenopausal and nulliparous, and (iv) postmenopausal and with at least 1 live birth.

Each donor underwent six core biopsies from the upper outer quadrant of one breast under local anesthesia. Within five minutes of collection, one biopsy was immediately placed into an embedding cassette and stored in 10% buffered formalin at room temperature before being embedded in paraffin. The remaining five biopsies were flash frozen in liquid nitrogen, then placed in labeled, chilled cryovials, and stored in liquid nitrogen, as described in our previous study^3,19^.

### DNA and RNA extraction

Breast tissue samples, each weighing 50 milligrams, were shipped to the Neurogenetics Core Sequencing Laboratory (UNGC) at UCLA for DNA methylation and transcriptome profiling. Frozen tissue (30 mg) was lysed in 600 μL of guanidine-isothiocyanate-containing Buffer RLT Plus in a 2.0 mL micro centrifuge tube and homogenized using TissueLyser II (Qiagen) with 5 mm stainless steel beads. The tissue lysate was then processed following the AllPrep protocol (Qiagen, catalog no. 80224) for simultaneous extraction of genomic DNA and total RNA, utilizing RNeasy Mini spin column technology. Extracted DNA underwent bisulfite conversion for methylation quantification, while the RNA was used for transcriptome quantification and analyses.

### DNA Methylation quantification and processing

DNA Methylation for each sample was measured using the Human Methylation EPIC (850K) array BeadChip (Illumina). 500 nanograms of DNA was bisulfite-converted using the EZ Methylation Kit (Zymo Research). Following bisulfite conversion, the DNA was hybridized to the EPIC array probes. Fluorescence data from the hybridized chip were scanned on an iScan (Illumina), where the methylated intensity (M) and unmethylated intensity (U) for each CpG site were measured.

Probe quality control and data processing were conducted using R package minfi (version 1.48.0). Specifically, we employed the ‘processIllumina()’ function to perform background subtraction and control normalization, and calculated DNA methylation level (beta-value) for each CpG site based on the intensity ratio between methylated (M) and an unmethylated (U) signal using the following formula

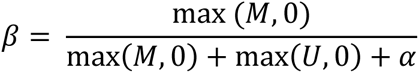

where α (default 100) is an offset to regularize beta value when both methylated and unmethylated probe intensities are low. By definition, the beta values range between 0 and 1, with 0 indicating completely unmethylated, and values approaching 1 indicating completely methylated.

### Bulk RNA sequencing and processing

Transcriptome profiling was performed using the Lexogen QuantSeq 3’ mRNA-Seq FWD kit to generate RNA sequencing libraries. Sequencing was conducted with 65 bp single end reads on an Illumina HiSeq 4000. The raw sequencing data underwent quality control using FastQC^49^ (version 0.11.9). Adapters and low-quality bases were trimmed using fastp^50^ (version 0.23.2). Trimmed reads were subsequently aligned to the human reference genome (GRCh38) with Ensembl annotation file (v84) using STAR^51^ (version 2.7.9a). Gene expression counts for each sample were obtained using HTSeq^52^ (version 1.99.2) and merged into a gene expression count matrix using in-house scripts. Genes with no more than 10 counts in less than 10% of samples were excluded from further analysis. Finally, the gene expression count per million (CPM) values were calculated using the ‘cpm()’ function implemented in the edgeR package^53^ (version 3.33.5).

### Sample quality control

Principal component analysis (PCA) was conducted on both the DNA methylation matrix (beta values) and the gene expression matrix (log-transformed CPM values) to identify potential sample outliers. After removing outliers and excluding samples from participants who developed breast cancer, we retained 181 normal breast tissue samples with paired DNA methylation and gene expression data. All downstream analyses were based on these 181 samples. The demographic and clinical characteristics of the finalized study cohort are detailed in Table 1 and Supplementary Data 1.

**Table 1.** Characteristics of the study population. SD: Standard deviation; n: number of samples; %: percentage; p-value: Two-tailed Student’s t-test for continuous variables and chi-square test for categorical variables. All participants in this analysis were of white race.

|  | Age < 50 years<br>(n=89) | Age >= 50 years<br>(n=92) | p-value |
| --- | --- | --- | --- |
| Demographics |  |  |  |
| Age (years), mean ± SD | 40.8 ± 7.7 | 59.0 ± 6.9 | <2.2e-16 |
| Ethnicity Hispanic, n (%) | 3 (3.4) | 6 (6.5) | 0.53 |
| Body mass index, mean ± SD | 28.2 ± 7.4 | 28.3 ± 6.2 | 0.93 |
| Tobacco smoking, ever, n (%) | 21 (23.6) | 34 (37) | 0.07 |
| Alcohol use, current, n (%) | 61 (68.5) | 64 (69.6) | 1 |
| Gynecologic history |  |  |  |
| Age at menarche, mean ± SD | 12.8 ± 1.6 | 12.7 ± 1.5 | 0.77 |
| Premenopausal, n (%) | 81 (91) | 14 (15.2) | <2.2e-16 |
| Age at menopause, mean ± SD | 36.7 ± 7.4 | 47.5 ± 6.6 | 0.0077 |
| Total menstrual years, mean ± SD | 27.5 ± 8.1 | 35.6 ± 6.5 | 8.9e-12 |
| Reproductive history |  |  |  |
| Nulliparous, n (%) | 43 (48.3) | 46 (50) | 0.94 |
| Age at first full-term birth, mean ± SD | 27.3 ± 5.3 | 26.0 ± 4.8 | 0.23 |
| Hormonal replacement therapy, ever, n (%) | 1 (1.1) | 37 (40.2) | <2.2e-16 |
| Estimated breast cancer risk scores |  |  |  |
| Gail score, mean ± SD | 13.3 ± 5.7 | 10.7 ± 5.3 | 0.0027 |
| Tyrer-Cuzick, lifetime, mean ± SD | 2.6 ± 1.9 | 4.4 ± 3.0 | 2.7e-06 |
| Tyrer-Cuzick, 10-year, mean ± SD | 15.6 ± 7.1 | 11.2 ± 6.4 | 2.5e-05 |

**Table 2.**
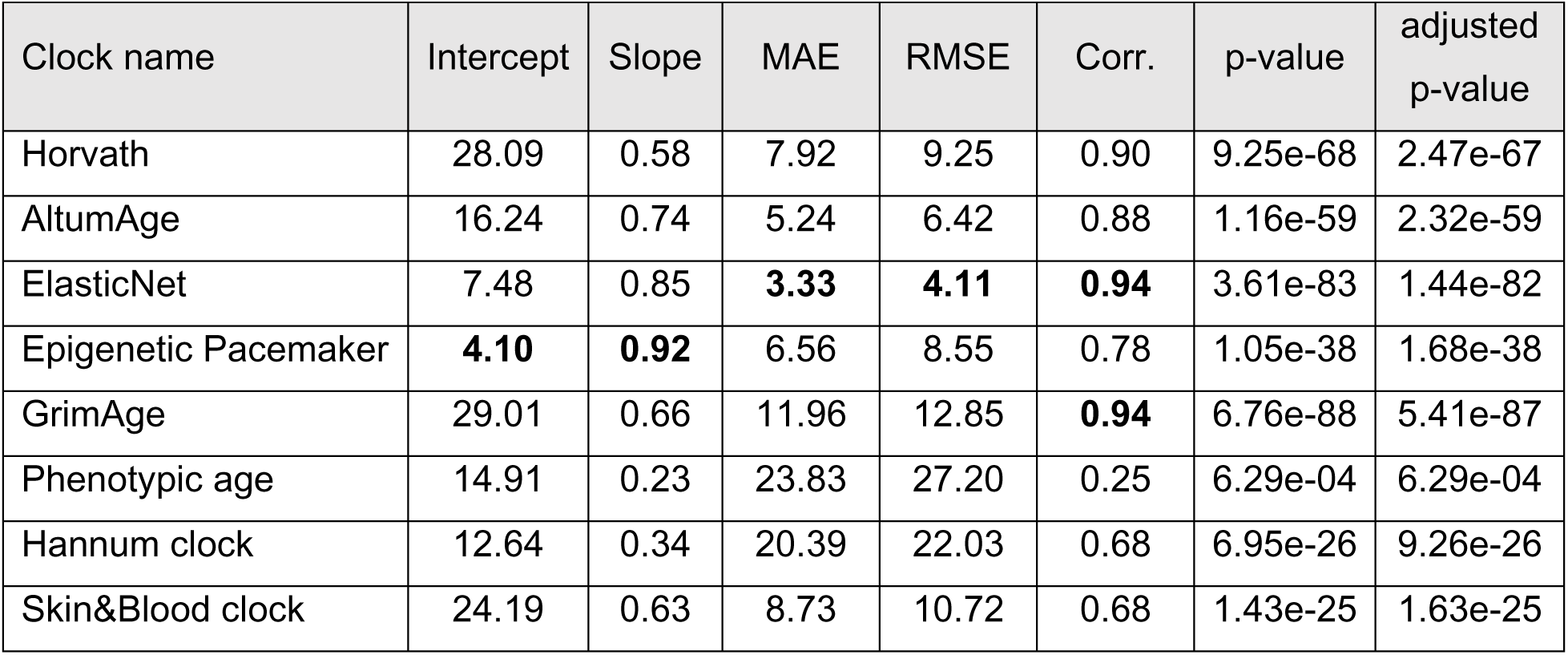
Regression summary for eight epigenetic clocks in KTB data. Intercept and Slope: the regression coefficients of epigenetic age against chronological age for each clock; MAE: Mean Absolute Error; RMSE: Root Mean Square Error; Corr.: Pearson Correlation coefficient; p-value: significance level of correlation via t-test; adjusted p-value: significance level of correlation adjusted by Benjamini-Hochberg procedure. The best prediction performance metrics (Intercept closest to 0, Slope closest to 1, smallest MAE/RMSE, and biggest Corr.) are highlighted in bold font.

### Epigenetic age and age acceleration calculation

To comprehensively assess the age prediction accuracy of DNA methylation (DNAm) epigenetic clocks in normal breast tissue, we included two pan-tissue clocks: Horvath’s pan-tissue clock^10^ and AltumAge^16^. These clocks are designed to operate across various tissue and cell types, including breast. Additionally, we examined two second-generation clocks, GrimAge^14^ and Phenotypic Age^12^, tailored to predict overall health span and lifespan more effectively, particularly in blood samples. GrimAge is of particular interest due to its potential to predict cancer onset and its association with menopausal age. Our analysis also included two first-generation clocks, the Hannum clock^11^ and the Skin&Blood Clock^13^, representing previous efforts for biological age prediction. The methodological details and applications of these clocks are further illustrated in Figure 1C, providing a clear summary of their distinct characteristics and diverse biological contexts. DNA methylation beta values were used to calculate the epigenetic age. Epigenetic age for Horvath, Hannum, GrimAge, Phenotypic Age, Skin&Blood was calculated using DNA methylation Age Calculator (https://dnamage.genetics.ucla.edu/home). AltumAge was calculated according to its official tutorial (https://github.com/rsinghlab/AltumAge).

Additionally, we trained two breast-specific clocks using the KTB DNA methylation data. One employs the Elastic Net algorithm^54^ to predict age based on methylation levels and the other utilizes Epigenetic Pacemaker (EPM) model^15^, which uses inverse regression to derive age estimates from DNA methylation patterns. To avoided data leakage, we implemented a nested cross-validation strategy: The outer loop used a leave-one-out approach to split the data into training and testing sets, while the inner loop employed a 5-fold cross-validation to tune the hyperparameters and train the epigenetic clocks. This strategy ensured that test data was never seen during model training. The trained model was then used to predict the age of each test data point, iterating through each sample to collect age predictions for all samples.

The relationship between chronological age and epigenetic age of different clocks was assessed using a generalized additive model with a cubic spline, we confirmed that the trend is predominantly linear on our data (Supplementary Figure 2A) and thus compute age acceleration by obtaining the residuals from a linear regression of epigenetic age on chronological age. This residual measure, designed to be age-adjusted, showed no correlation with chronological age (Supplementary Figure 3B). Despite concerns about sampling uncertainty affecting the residual calculations, we used bootstrapping to confirm that age acceleration of each subject remains stable against variations in sample composition for each clock (Supplementary Figure 3C). The robust measurement underscores the reliability of age acceleration and our findings.

### Cell type deconvolution and immune enrichment score calculation

To quantify the cell type abundance in normal breast tissue samples, we utilized the Genotype-Tissue Expression (GTEx) v8 single-nucleus RNA-seq (snRNA-seq) data (https://gtexportal.org/home/downloads/single_cell) and extracted gene expression data specific to normal breast tissue. The snRNA-seq breast dataset identified eight major cell types: adipocytes, luminal epithelial cells, basal myoepithelial cells, vascular endothelial cells, lymphatic endothelial cells, immune cells (dendritic cells/macrophages), pericytes & smooth muscle cells, and fibroblasts (Supplementary Figure 4A, 4B). Cell doublets and genes expressed in fewer than 10 cells were excluded from the snRNA-seq data. The processed expression count matrix was then converted into counts per million (cpm) values. Using this data as a reference, we constructed a cell type signature matrix and deconvolved the bulk RNA-seq data to determine cell type abundance proportions for each sample using CIBERSORTx^30^ with batch correction mode (S-mode).

To achieve higher resolution of immune cell composition in normal breast tissue, we used ImmuneCellAI^32^ (http://bioinfo.life.hust.edu.cn/ImmuCellAI/) to calculate enrichment scores for 24 immune cell types and immune infiltration scores for each sample based on gene expression data. All software parameters were set as default unless otherwise specified.

### Cancer risk score calculation

In addition to the Gail and Tyrer-Cuzick breast cancer risk measurements, which were computed in previous work using demographic, reproductive and family history data^3^, we expanded our analysis to include cancer risk scores derived from molecular data in this analysis. Specifically, we utilized the code provided in epiTOC2^34^ to compute cancer risk scores for each sample based on the DNA methylation matrix. We also calculated cell senescence scores using gene expression data by applying single-sample Gene Set Enrichment Analysis (ssGSEA) with senescence signature genes from the CellAge database^55^, implemented through the GSVA package^56^ (version 1.50.5). By integrating these molecular-based risk scores with traditional risk assessments, we aimed to provide a more holistic view of breast cancer risk. The processed data, including epigenetic age estimates, age acceleration of each clock, cell proportions, immune enrichment scores, and cancer risk scores can be found in Supplementary Data 2.

### Correlation and Mediation analysis

We computed the pairwise Pearson Correlation Coefficients among various variables of interest. These include demographic variables, reproductive history, breast cancer risk and senescence scores, epigenetic age from eight different clocks, age acceleration (both raw and adjusted by cell proportions), the top 10 principal components from DNA methylation and gene expression, eight cell type proportions, 24 immune cell scores, and immune infiltration scores. The comprehensive correlation matrix is provided in Supplementary Data 3 and visualized in Supplementary Figure 6.

To examine how changes in cell type proportions mediate the relationship between chronological age and epigenetic age, we conducted a mediation analysis. In this analysis, chronological age served as the independent variable (IV), epigenetic age as the dependent variable (DV), and each cell type abundance as mediators. The structural equation model used for this analysis is detailed in the Supplementary Data 4. We utilized the R package lavaan^57^ (version 0.6-18) to perform the mediation analysis for each clock, estimating both direct and indirect effects and the contribution of each mediator.

### Epigenome-wide association study (EWAS) and genomic region enrichment analysis

We conducted an Epigenome-Wide Association Study (EWAS) to identify CpG sites associated with age acceleration, controlling for chronological age and cell proportions. The EWAS analysis was performed using GEMMA^58^ (version 0.98.6), where the age acceleration for each clock is the quantitative traits and the CpG sites are the markers, with age and cell proportions as covariates. Since the cell proportions sums up to 1, we excluded Pericyte/SMC from the covariates to avoid model identifiability issue. The association between each CpG site and age acceleration was then tested using a linear mixed model framework in GEMMA, with p-values calculated via likelihood ratio tests. CpG sites with q-values smaller than 0.05 were considered significantly associated (Supplementary Data 5) and were subsequently subjected to genomic region enrichment analysis using LOLA^59^ (version 1.32.0), with all CpG sites serving as the background (Supplementary Data 6). The top 10 most significantly enriched Transcription Factor Binding Site (TFBS) were extracted and visualized in Figure 5.

### Differential expression and gene-set enrichment analysis

To further explore the relationship between gene expression and age acceleration for each clock, we conducted differential expression (DE) analysis. First, we converted the gene expression matrix (CPM) for each gene by applying an inverse Gaussian transformation, ensuring that each gene’s expression value *Y_i_* follows a Gaussian distribution. Next, we performed DE analysis using a linear regression framework with age and age acceleration (AA) as covariates, both before (1) and after (2) adjusting for cell proportions:

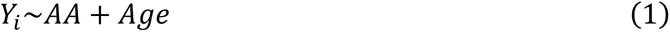

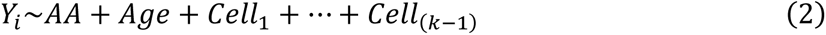

Where AA is the age acceleration of a specific clock, and k=8 is the total number of major cell types. Since the cell proportions sums up to 1, we excluded Pericyte/SMC when fitting the second model to avoid model identifiability issue. We implemented these models using ‘lm()’ function in R. For both models, we tested whether AA is associated with the response *Y_i_* for each gene. The association p-values were obtained using t-tests (Supplementary Data 7).

Genes with an adjusted p-value less than 0.05 in model (2) were defined as differentially expressed for each clock and subjected to Gene Ontology (GO) overrepresentation enrichment analysis. Additionally, we performed Gene Set Enrichment Analysis (GSEA) using the differential analysis results. Both GO overrepresentation and GSEA analyses were conducted and visualized using clusterProfiler^60^ (version 4.10.1). The gene expression enrichment analysis results is provided in Supplementary Data 8.

### Statistical analysis

Besides the statistical analysis described above, the pairwise Pearson Correlation Coefficient and significance (p-values) are calculated using R package Hmisc^61^ (version 5.1-2). Correlation heatmaps were used to display the pearson correlation coefficients using R package corrplot^62^ (version 0.92). To account for multiple hypothesis testing, we reported the adjusted p-values using Benjamini-Hochberg^63^ procedure implemented by ‘p.adjust()’ function in R. All statistical tests were conducted in R (version 4.3).

## Data availability

Raw data will be available upon manuscript publication. Processed data is provided in the Supplementary Data 1 and 2.

## Acknowledgement

We thank the study donors and contributors to the Susan G. Komen Tissue Bank at the IU Simon Cancer Center whose participation, data collection, and support made this work possible. This study was funded by a Susan G. Komen Career Catalyst Award for Basic and Translational Science CCR16380478. W.G. was funded in part by the UCLA QCBio Collaboratory Fellowship. P.A.G was funded in part by the Breast Cancer Research Foundation. Figure 1A was created with BioRender.com, thus we acknowledge it by courtesy.

## Author contributions

M.E.S., W.G., S.H., M.P., and P.A.G. contributed to the conceptual design, data curation, resources, and writing of the manuscript. W.G. performed all the analyses and drafted the manuscript with assistance from M.E.S. J.P. contributed to the data curation and manuscript editing. C.F. and J.J.L. contributed to the data analysis and manuscript editing. A.M.S., N. M., and J.E.H. contributed to the resources, data curation, manuscript editing. M.E.S, M.P, P.A.G. jointly supervised the study.

## Competing interests

P. A. G is a member of the foundation scientific advisory board. S.H. is a founder of the nonprofit Epigenetic Clock Development Foundation, which has licensed several patents from UC Regents, and distributes the mammalian methylation array. M.P. founded ProsperK9.

## List of Tables and Figures

**Supplementary Figure 1.**
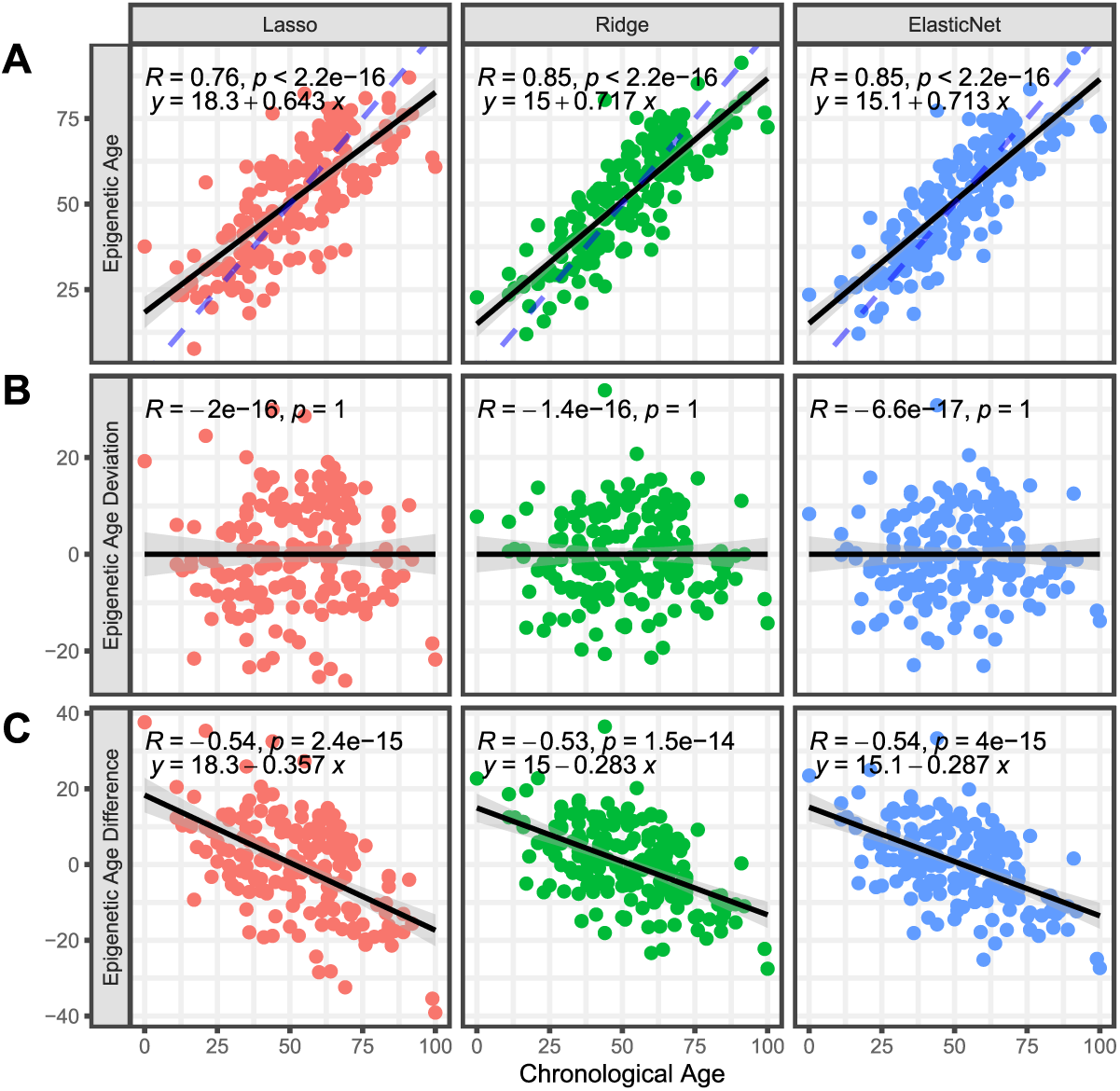
Age deviation and Age difference comparison across different penalized regression methods in simulation. Scatter plot shows chronological age is strongly correlated with predicted/epigenetic age (**A**), not correlated with Age deviation (**B**), and inversely correlated with Age difference (**C**) across penalization methods in simulation, indicating the confounding issue of using difference as age acceleration measurement. Columns represent the three different penalization techniques (Lasso: least absolute shrinkage and selection operator, Ridge: Ridge regression, ElasticNet: Elastic Net regression). Black line represents the regression line with shadowed region indicating 95% Confidence Interval.

**Supplementary Figure 2.**
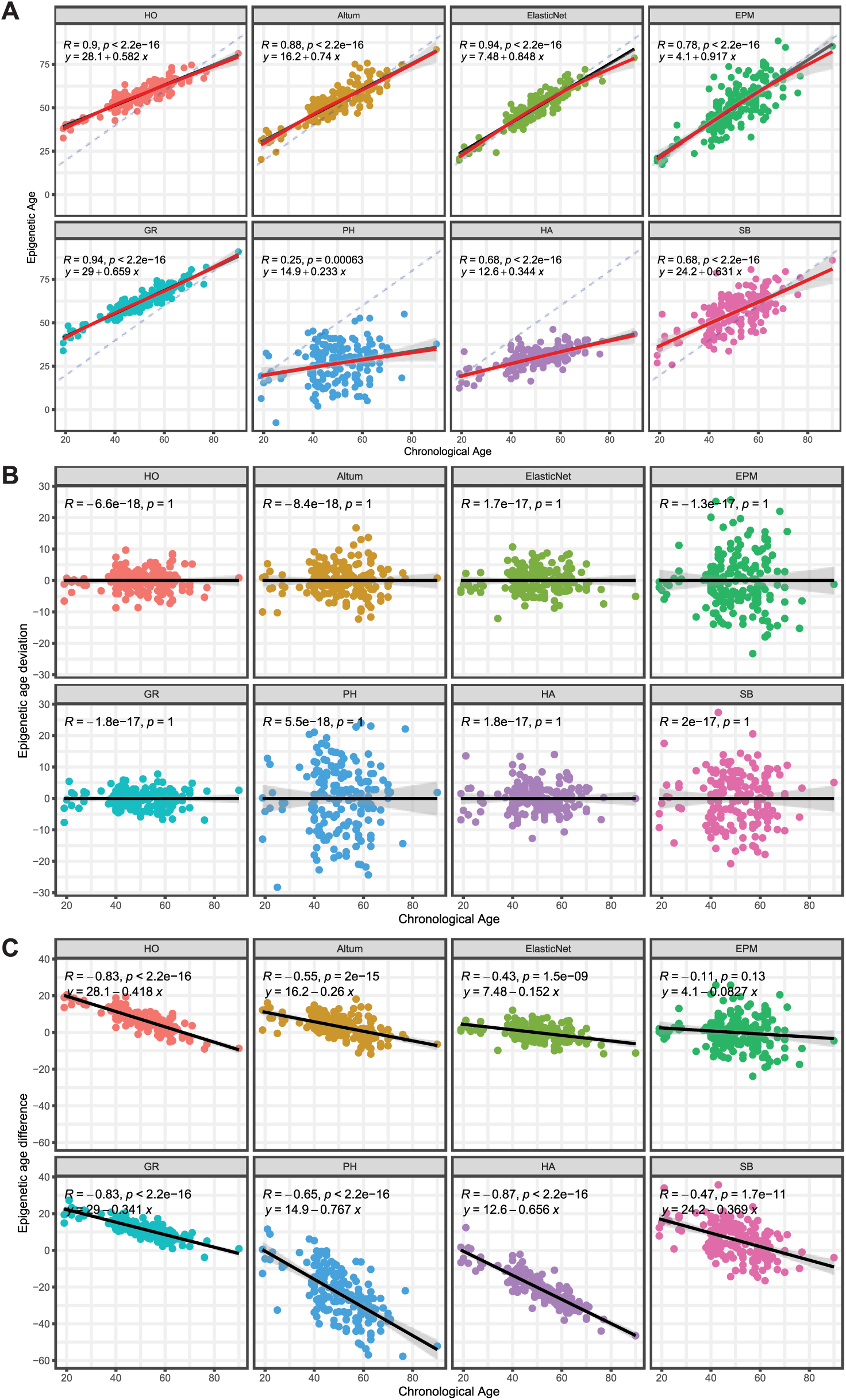
Age deviation and Age difference comparison across epigenetic clocks in KTB data. Scatter plot shows chronological age is strongly correlated with predicted/epigenetic age (**A**) in KTB data, with redline representing the cubic spline regression and confirming the change tendency between epigenetic age and chronological age is predominantly linear across epigenetic clocks. Chronological age is not correlated with Age deviation (**B**), and inversely correlated with Age difference (**C**) in real data, indicating the confounding issue of using difference as age acceleration measurement. Black line represents the regression line with shadowed region indicating 95% Confidence Interval.

**Supplementary Figure 3.**
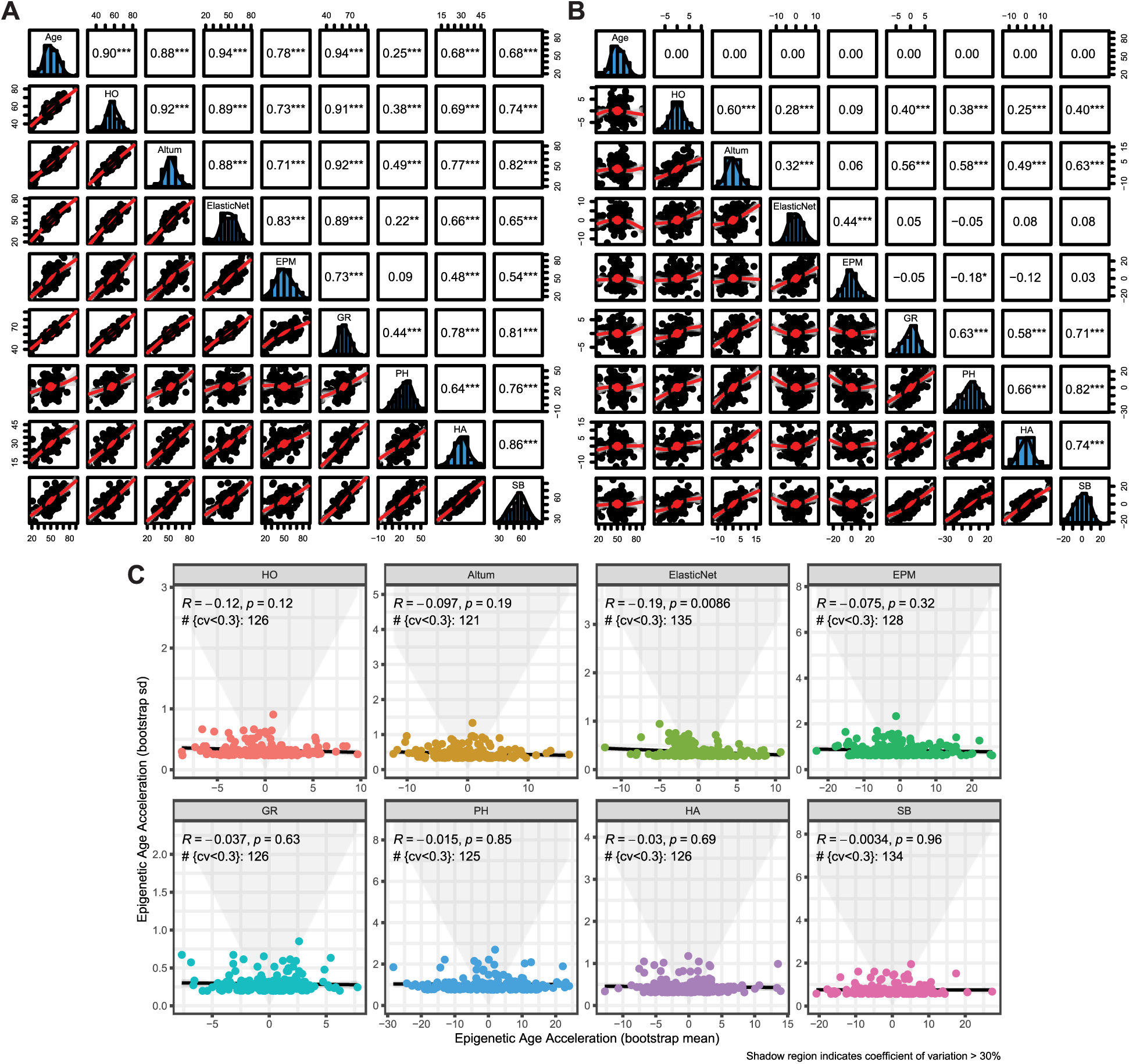
Correlations among epigenetic age, age acceleration, and the robustness of age acceleration to sampling in KTB data. (**A**-**B**) Scatterplot matrices for epigenetic age (**A**) and age acceleration (**B**), with chronological age added as a reference. Lower panels show the scatter plot of a variable pair. Upper panels display the pairwise Pearson Correlation Coefficient and significance level (*: p < 0.05, **: p< 0.01, ***: p < 0.001)**. (C)** Robustness of age acceleration to variations of sample composition. Bootstrap sampling was conducted for 1000 times with bootstrap mean (x-axis) and standard deviation (y-axis) of age acceleration calculated for each sample, shadowed region represents coefficients of variation (cv) greater than 0.3, showing most samples have stable age acceleration across clocks.

**Supplementary Figure 4.**
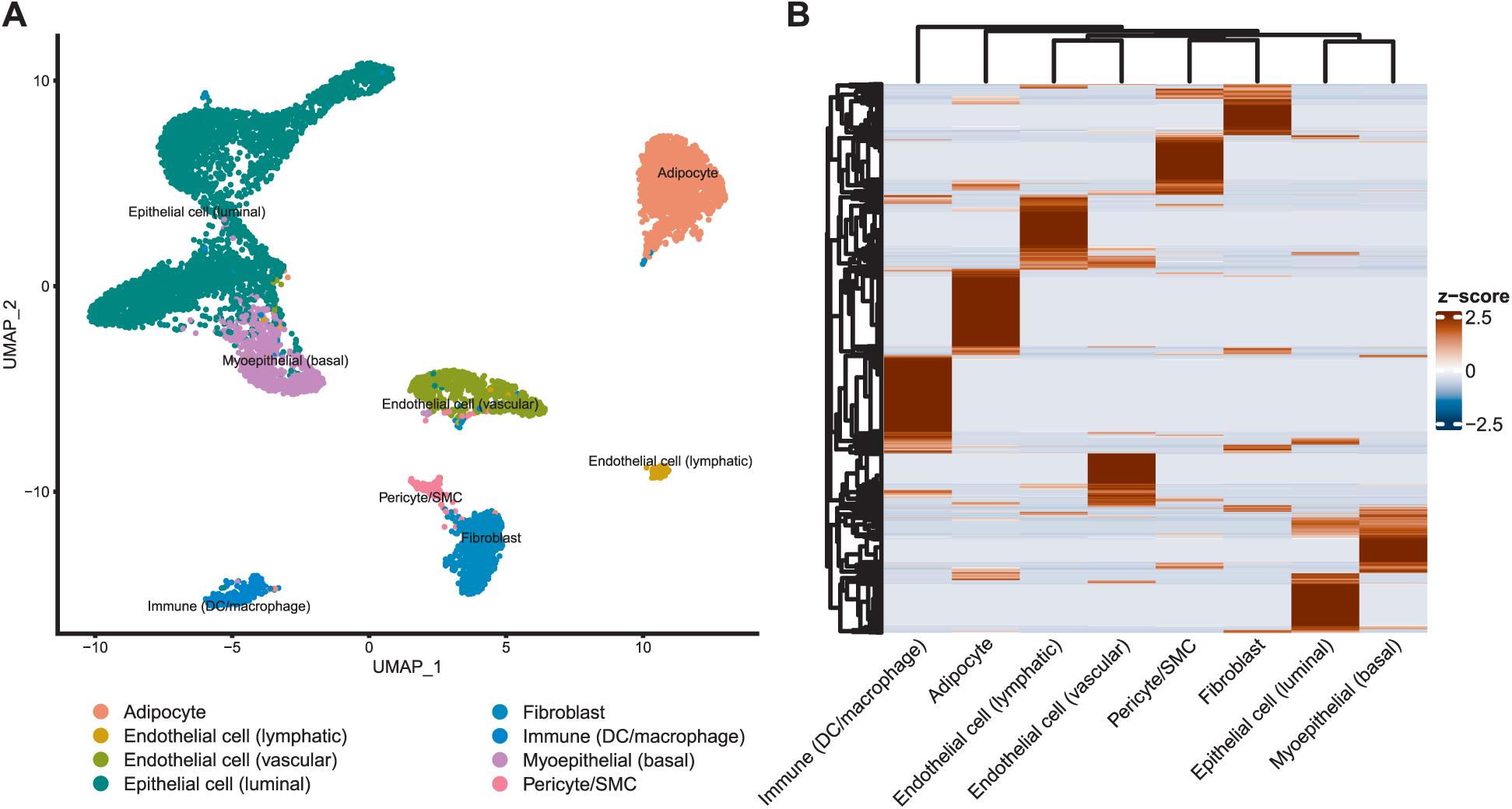
GTEx normal breast snRNA data for cell deconvolution. **(A)** UMAP showing eight major cell types in GTEx normal breast tissue snRNA-seq data. **(B)** Heatmap showing the single cell gene signature matrix constructed by CIBERSORTx and used for cell deconvolution, the standardized gene expression value (z-score) is shown for each cell type.

**Supplementary Figure 5.**
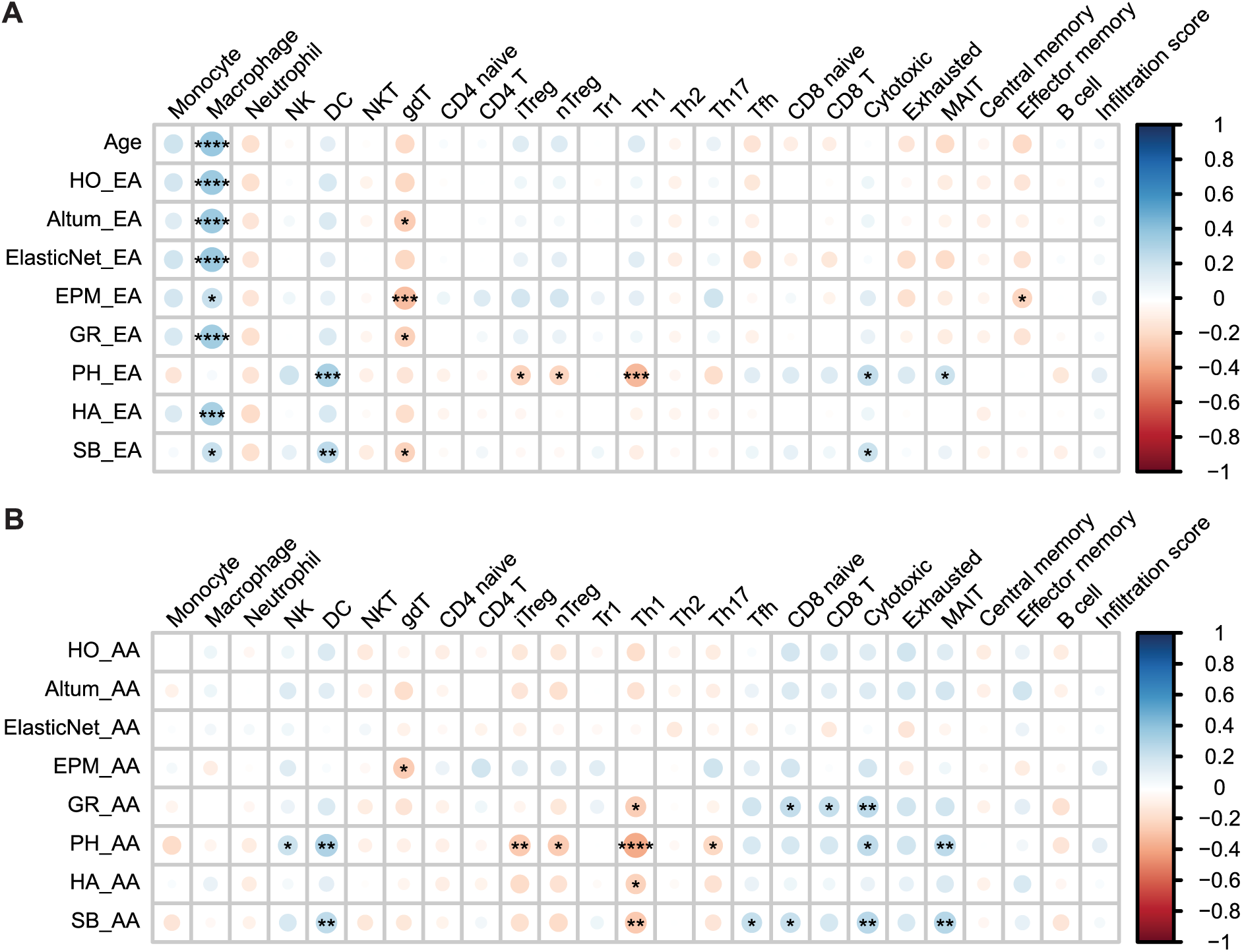
Association of immune cell scores with epigenetic age and age acceleration. Correlation heatmap showing the relationship between immune cell scores and epigenetic age (**A**) and age acceleration (**B**) across epigenetic clocks. Chronological age was added to the top row as a reference. Circle size and color scale representing the strength of correlation with blue color shows positive correlation and red color indicates negative correlation. Asterisk indicates significance level after p-value adjustment. *: adjusted p-value < 0.05; **: adjusted p-value < 0.01; ***: adjusted p-value < 0.001; ****: adjusted p-value < 0.0001.

**Supplementary Figure 6.**
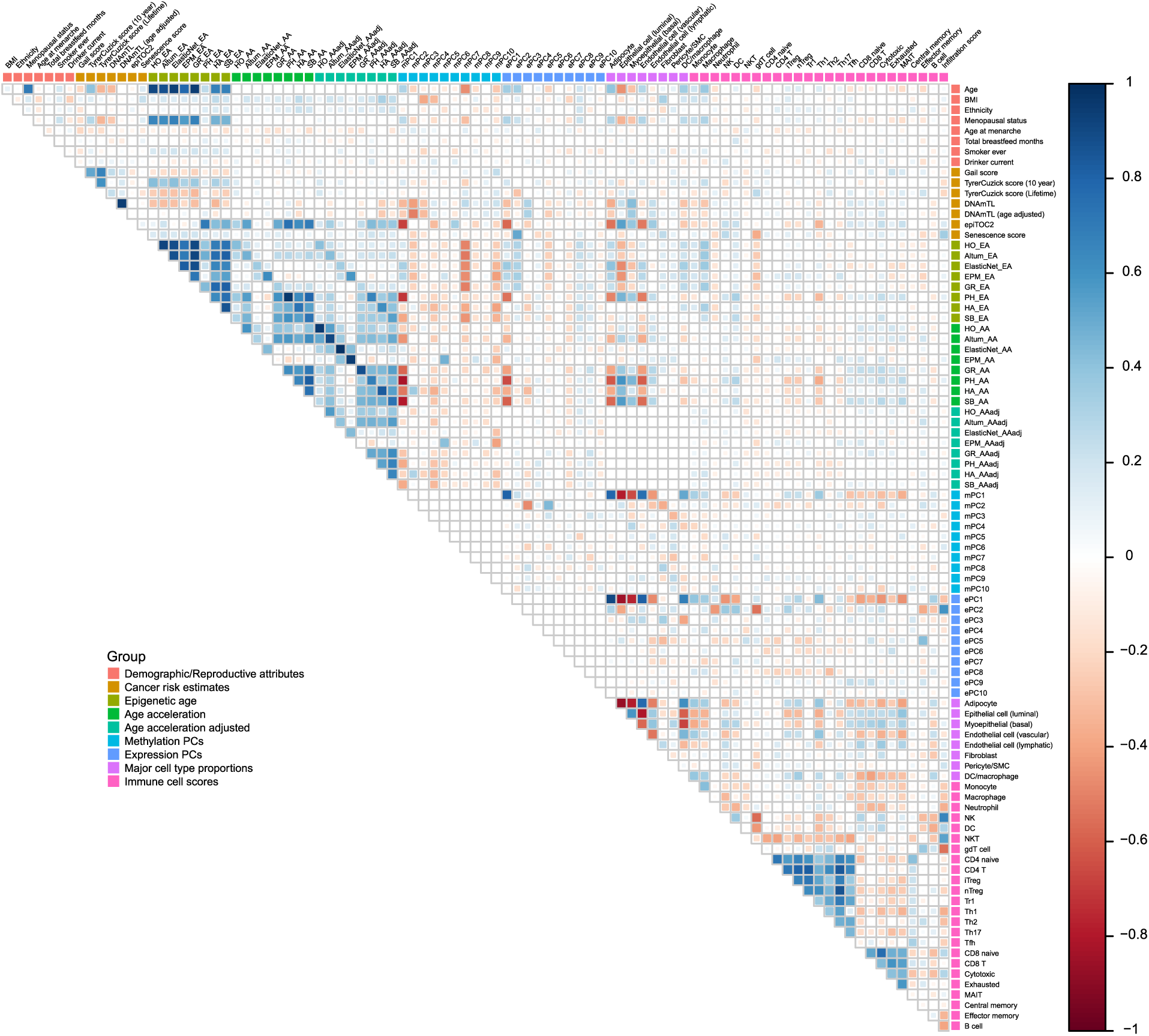
Pairwise correlation heatmaps of examined variables. Correlation heatmap showing the pairwise correlation among demographical variables (Age, Ethnicity, BMI, Smoker ever, Acholic drinks per week), reproductive history (Menopausal status, Age at menarche, Total breastfeed months), breast cancer risk estimates, eight epigenetic clocks’ epigenetic age, age acceleration, age acceleration adjusted by cell proportions, top 10 principal components from DNA methylation (mPC1-mPC10) and gene expression (ePC1-ePC10), eight cell proportions, 24 immune cell scores and immune infiltration score.

**Supplementary Figure 7.**
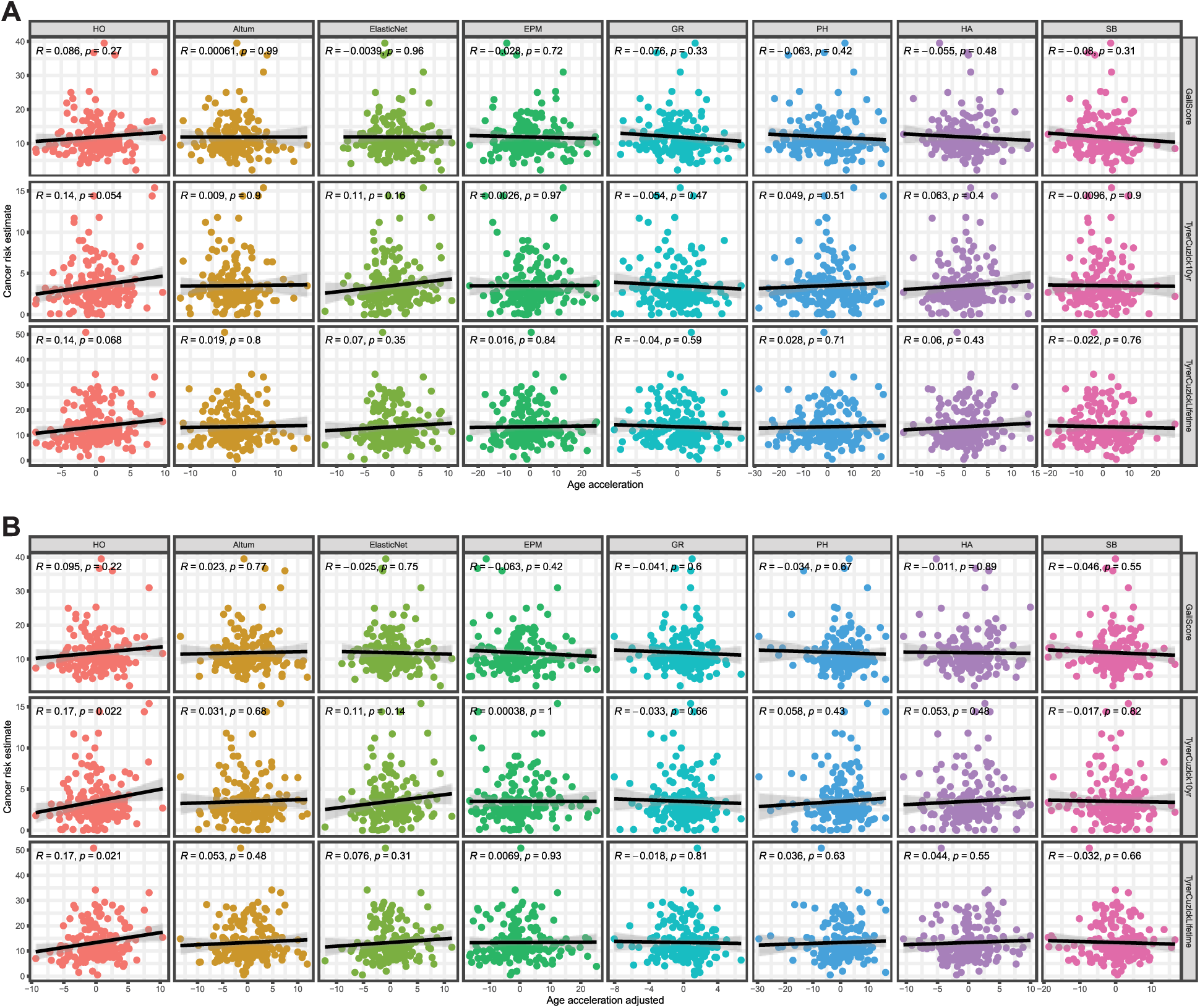
Association between cancer risk estimates and age acceleration. Scatterplot showing the association between cancer risk measurement and age acceleration before (**A**) and after (**B**) adjusting cell proportions. Columns show the eight epigenetic clocks and rows represent the three breast cancer risk measurements. Black line represents the regression line with shadowed region indicating 95% Confidence Interval. Pearson Correlation Coefficient statistics are annotated on the top of each panel.

**Supplementary Figure 8.**
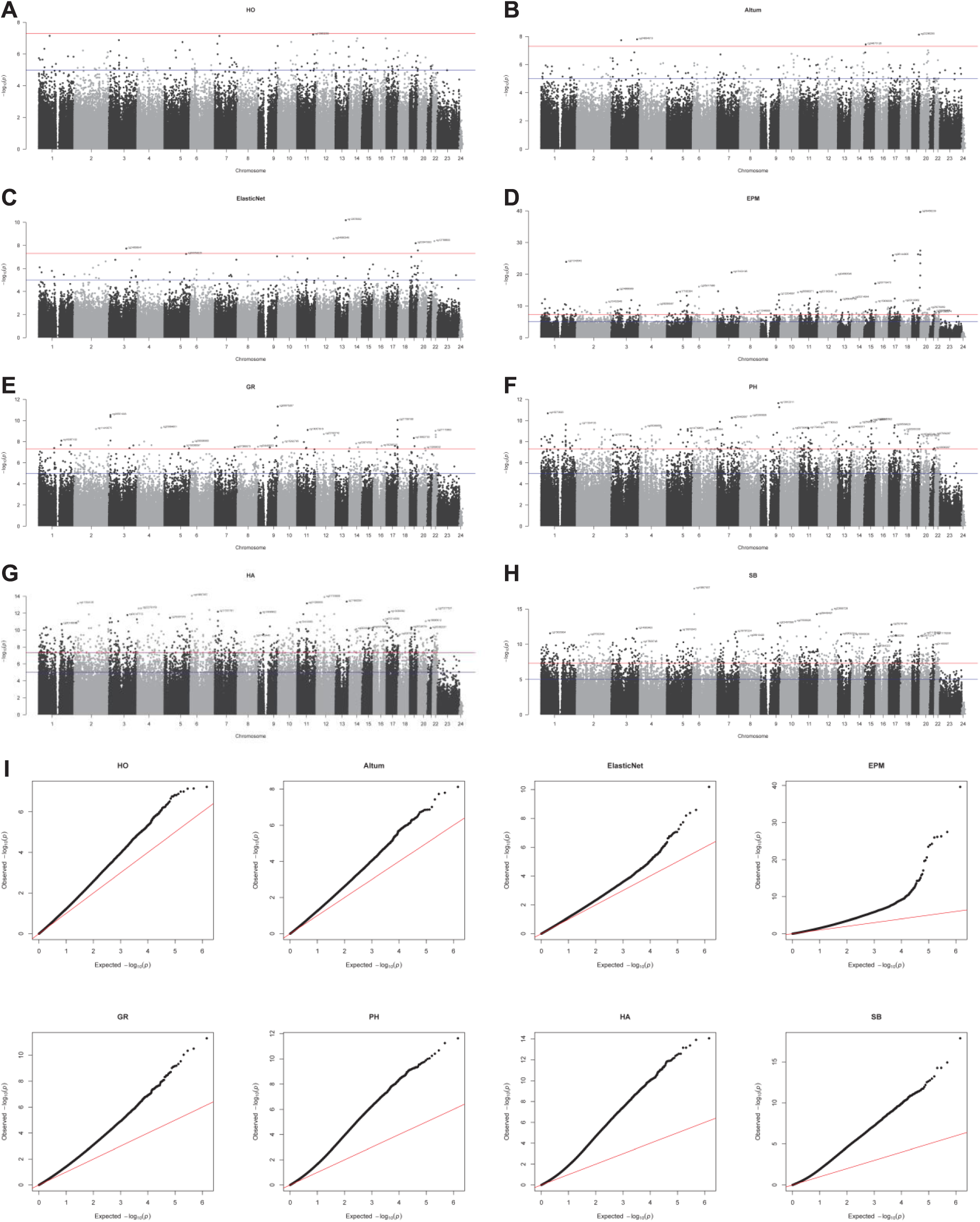
Manhattan and QQ plot for EWAS analysis. (**A**-**H**) Manhattan plot of EWAS analysis for pan-tissue clocks: Horvath clocks (**A**) and AltumAge (**B**), breast-specific clocks: clock trained using ElasticNet algorithm (**C**) and Epigenetic pacemaker (**D**), second generation clocks: GrimAge (**E**) and Phenotypic age (**F**), first generation clocks Hannum clock (**G**) and Skin&Blood clock (**H**); Blue and red horizontal line on the Manhattan plot shows 10^-5^ and 5*10^-8^ p-value threshold, respectively. Top sites on each chromosome whose p-value passed Bonferroni correction threshold are labeled onto the plot. (**I**): Quantile-Quantile (QQ) plot for all eight clocks examined.

**Supplementary Figure 9.**
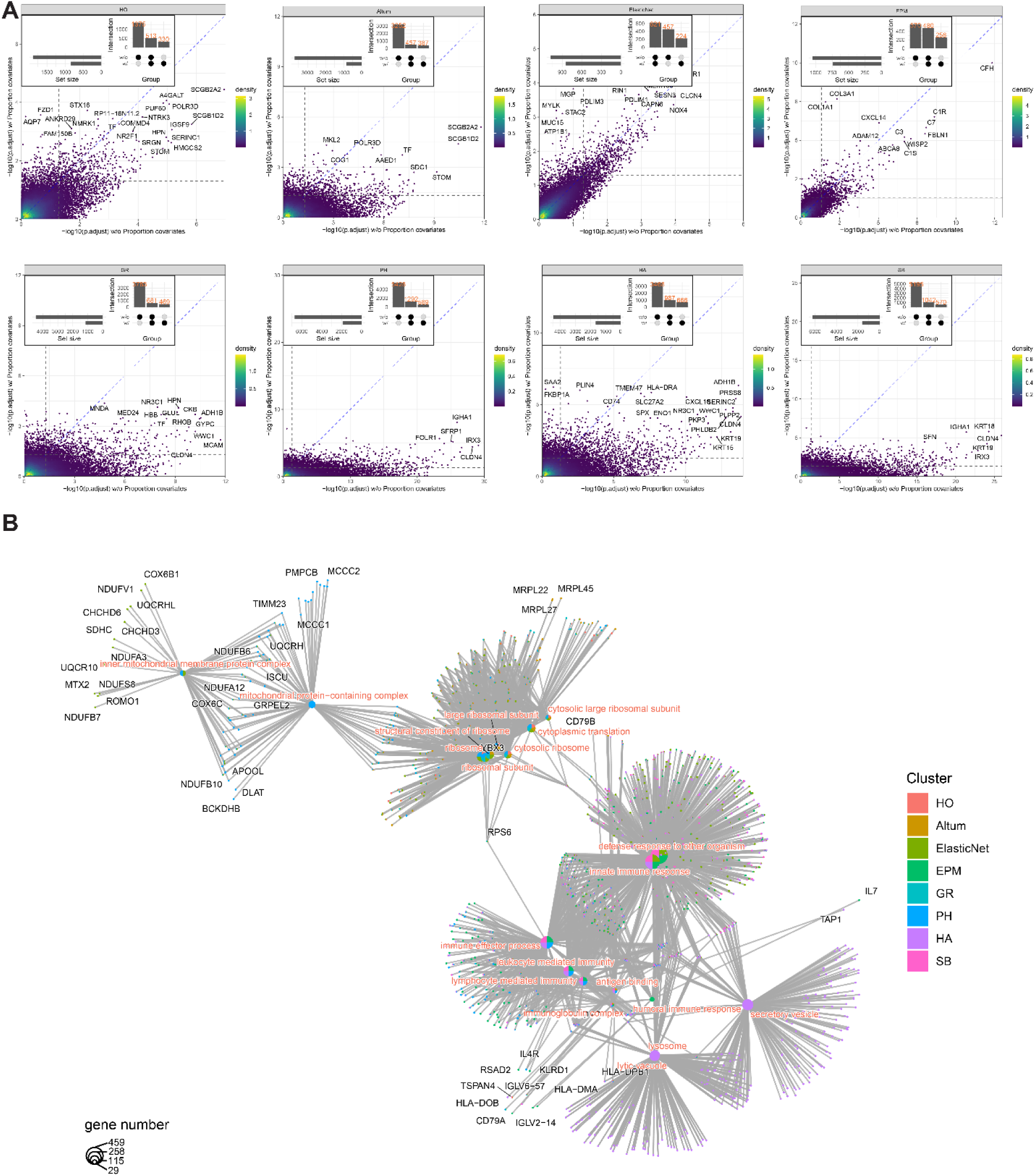
Differential gene expression and gene set enrichment analysis. Differential expression (DE) analysis was performed to identify genes associated with age acceleration for each clock, followed by gene set enrichment analysis. **(A)** scatterplot showing the DE analysis before and after adjusting cell proportions. Each dot is a gene with -log10 of adjusted p-value before (x axis) and after (y axis) adjusting cell proportions. Vertical and horizontal dashed line indicates 0.05 threshold. Genes with adjusted p-value smaller than 0.05 are defined as differential expressed (DE) genes, and the number of DE pre/post adjusting cell composition as well as their overlap were summarized using the bar plot annotated to the top-left of the scatter plot. The genes accumulating below the diagonal dashed line indicates many of the differential genes are due to cell compositional changes. **(B)** Cnetplot displaying the GSEA pathway enrichment analysis results. Large nodes with pinkish font represent pathways, while small nodes with black font represent genes connected to the pathways by edges. Node size represents the number of genes of a pathway and the colors of large nodes represent whether the given pathway is enriched in the corresponding clocks.

